# Arginine vasopressin activates serotonergic neurons in the dorsal raphe nucleus during neonatal development *in vitro* and *in vivo*

**DOI:** 10.1101/2024.03.22.586208

**Authors:** Ester Orav, Bojana Kokinovic, Heidi Teppola, Mari Siimon, Sari E. Lauri, Henrike Hartung

## Abstract

Birth stress is a strong risk factor for psychiatric disorders and associated with an exaggerated release of the stress hormone arginine vasopressin (AVP) into circulation and in the brain. While it has been shown that AVP promotes firing of GABAergic interneurons leading to suppression of spontaneous perinatal hippocampal network events that suggest a protective function, its effect on developing subcortical networks is not known. Here we tested the effect of AVP on the neonatal dorsal raphe nucleus (DRN) 5-hydroxytryptamine (5-HT, serotonin) system, since early 5-HT homeostasis is critical for the development of cortical brain regions and emotional behaviors. Using *in vitro* electrophysiological recording techniques, we show that AVP strongly excites neonatal 5-HT neurons via V_1A_ receptors by increasing their excitatory synaptic inputs. Accordingly, AVP also promotes action potential firing through a combination of its effect on glutamatergic synaptic transmission and a direct effect on the excitability of 5-HT neurons. Our *in vivo* single unit recordings of identified neonatal 5-HT neurons under light urethane anaesthesia revealed two major firing patterns of neonatal 5-HT neurons, tonic regular firing and low frequency oscillations of regular spike trains. We confirmed that AVP also increases firing activity of putative 5-HT neurons in neonatal DRN *in vivo*. Finally, we show that neonatal DRN contains a sparse vasopressinergic innervation that is strongly sex dependent and originates exclusively from vasopressinergic cell groups in medial amygdala and bed nucleus of stria terminalis (BNST). Our results show, that in contrast to developing cortical networks where AVP promotes inhibition, AVP can also be strongly excitatory in immature subcortical networks such as the DRN 5-HT system. Hyperactivation of the neonatal 5-HT system by AVP during birth stress may impact its own ongoing functional development as well as affect maturation of cortical target regions, which may increase the risk for psychiatric conditions later on.

**Author Contributions:** E.O. performed and analysed the *in vitro* electrophysiological experiments, related immunohistochemistry of filled neurons as well as image analysis, B.K. and H.H. conducted and analysed the *in vivo* juxtacellular electrophysiological recordings and labelling, related immunohistochemistry of labelled neurons and image analysis, H.H. did the multi-channel *in vivo* electrophysiological recordings and intracerebral injections as well as related histology, H.T-G. analysed the multi-channel *in vivo* electrophysiological data, B.K. and H.H. performed the tracing experiments, E.O. and H.H. carried out immunohistochemistry related to the tracing experiments, E.O. performed image analysis related to tracing experiments, M.S. performed and analysed AVP immunocytochemistry experiments in neonatal DRN, H.H. and S.E.L. provided resources for the experimental work and supervised the project. H.H. conceptualized and coordinated the project. The manuscript was written by H.H. with significant contributions from all authors.

## INTRODUCTION

There is an increasing consensus that many neuropsychiatric disorders have a developmental origin since early-life adverse events (ELAs) make individuals strongly susceptible to later disease (Kessler et al., 2010; McKay et al., 2021; McKay et al., 2022).

One of the early risk factors are obstetric complications including long and complicated labour or mild birth asphyxia that cause stress to the infant and make individuals more susceptible to psychiatric disorders such as affective disorders or schizophrenia (Nosarti et al., 2012; Schmitt et al., 2014; Buoli et al., 2016; Belbasis et al., 2018; Álvarez-García et al., 2022; Schmitt et al., 2023). Birth stress, in common with other ELAs such as neglect or abuse, leads to strong activation of the neonatal stress axis during a sensitive period of brain development. Normal vaginal birth is accompanied by a surge of the stress hormone arginine vasopressin (AVP) as well as catecholamines and cortisol into the blood stream of the fetus (Evers and Wellmann, 2016). Indeed, studies in rodents have shown that AVP neurons particularly in paraventricular nucleus of the hypothalamus (PVN) and supraoptic nucleus (SON) are activated during birth (Hoffiz et al., 2021) and release AVP via the posterior pituitary into circulation. AVP release is triggered by periods of hypoxia during labour contractions as shown in prenatal sheep (DeVane et al., 1982; Pimentel et al., 1989) and has also been observed in rodent models of birth asphyxia (Summanen et al., 2018). Accordingly, AVP levels in human cord blood are about 100-fold higher after vaginal delivery than after delivery by elective C-section (Evers and Wellmann, 2016). The strong release of stress hormones and catecholamines during birth serves to quickly adapt to extrauterine life. Their function of facilitating air breathing and the adaptation of blood flow and pressure, body temperature as well as glucose and water homeostasis promotes survival of the neonate (Hillman et al., 2012). During asphyxic birth the surge of AVP is even higher reaching approx. twice the levels during normal vaginal birth (Schlapbach et al., 2011). Interestingly, there is evidence that AVP concentrations in cerebrospinal fluid (CSF) of healthy neonates but also neonates with hypoxic-ischemic encephalopathy are very similar and correlate well with plasma levels (Bartrons et al., 1993; Carson et al., 2014). This suggests that the surge of AVP during birth occurs not only in the periphery but also in the brain via central projections of hypothalamic AVP neurons. Furthermore, while it is not known whether also extra-hypothalamic AVP neurons in the bed nucleus of the stria terminalis (BNST) and medial amygdala are directly activated during birth, recent anatomical data in mice show that AVP neurons in BNST and medial amygdala are highly interconnected and receive inputs from a substantial proportion of PVN AVP neurons (Rigney et al., 2023). This suggests a coordinated action which may also lead to activation of extra-hypothalamic AVP neurons during birth indirectly via activation of AVP neurons in PVN (Rigney et al., 2023). This combined activation may also contribute to increased AVP release during birth in the brain.

In line with its survival-promoting function in the periphery, it has been proposed that AVP protects the perinatal brain during birth by promoting inhibition of brain activity. Indeed, AVP at nanomolar concentrations suppresses spontaneous neuronal network activity in the perinatal hippocampus *in vitro* by increasing the firing activity of local GABAergic interneurons and inhibitory synaptic input onto pyramidal neurons (Spoljaric et al., 2017). However, if this early inhibitory function in the hippocampus is universal and also applies to other brain regions including subcortical networks is not known.

The neuromodulator 5-hydroxytryptamine (5-HT, serotonin) is a powerful regulator of emotional behaviors in adulthood and has long been implicated in psychiatric disorders such as depression and anxiety disorders. Furthermore, the development of emotional behaviors is also strongly dependent on early 5-HT homeostasis: a postnatal increase in 5-HT by pharmacologically blocking the serotonin transporter (SERT) results in depressive-like and anxiety behavioral traits in adult rodents (Ansorge et al., 2004; Ansorge et al., 2007; Gingrich et al., 2017).

Hence, while in adulthood decreased 5-HT levels in the brain seem to be associated with affective disorders, paradoxically, during neonatal development, abnormally increased levels also result in impaired emotional behaviors later on (Suri et al., 2015). The vast majority of 5-HT neurons projecting to the forebrain are located in the dorsal raphe nucleus (DRN). The DRN also receives a dense vasopressinergic innervation (Rood and De Vries, 2011; Rood and Beck, 2014) originating from vasopressinergic cell groups in BNST and medial amygdala (Rood et al., 2013; Rigney et al., 2023) and shows expression of vasopressin 1A (V_1A_) receptors (Ross et al., 2019; Patel et al., 2022). Moreover, both rodents and humans have a strong transient developmental expression of V_1A_ receptors in cortical and subcortical brain regions around the time of birth (Kang et al., 2011; Hammock and Levitt, 2012; Hammock, 2015). The function of AVP in DRN is still largely unknown. In adulthood, AVP has been shown to increase the excitatory synaptic input onto DRN 5-HT neurons via a V_1A_ – dependent glutamatergic mechanism (Rood and Beck, 2014). Moreover, 5-HT neurons in DRN show a protracted functional development that is characterized by early hyperexcitability during neonatal development before reaching maturity after the third postnatal week in mice (Rood et al., 2014).

We hypothesized that in contrast to its inhibitory effect in hippocampal or cortical networks, AVP would be highly excitatory in the immature DRN leading to increased firing of neonatal serotonergic neurons *in vitro* and *in vivo*. Since serotonin affects circuit development in target brain regions (Gaspar et al., 2003), increased and prolonged release of AVP during long and complicated labour may lead to strong activation of the neonatal 5-HT system and increased 5-HT levels that may contribute to the increased susceptibility to psychiatric disorders after birth stress.

We tested the response of developing 5-HT neurons in the rat DRN to AVP at postnatal day (P) 10-12, a time corresponding to human birth by electrophysiological recordings either *in vitro* in acute brain slices or *in vivo* in anaesthetized rat pups. We show that AVP at nanomolar concentrations strongly increases the excitatory synaptic drive to identified neonatal 5-HT neurons via V_1A_ receptors and leads to a strong increase in action potential firing *in vitro*. This effect is a combination of an indirect glutamatergic mechanism and a direct effect on 5-HT neurons. Our *in vivo* electrophysiological recordings provide the first characterization of the firing activity of identified neonatal 5-HT neurons *in vivo* and show that a large proportion of immature 5-HT neurons exhibit low-frequency oscillations in firing. Furthermore, we confirm that local AVP injections also increase the firing activity of putative 5-HT neurons *in vivo*. We reveal, that the neonatal DRN already displays a vasopressinergic innervation, which, despite its sparseness, already displays typical features of the adult innervation such as strong sex dependence. Retrograde tract tracing experiments confirm that the innervation emerges from cell groups in medial amygdala and BNST with approximately equal contributions.

Our results show that the inhibitory function of AVP, observed in cortical networks at birth is not universal throughout the brain and that AVP can also be strongly excitatory in subcortical networks such as the immature DRN. While a fine balance between inhibitory and excitatory effects of AVP in immature networks protects the brain during normal vaginal birth, prolonged and increased activation of the AVP system during complicated birth or mild birth asphyxia may cause this balance to break down. Prolonged AVP-mediated activation of the 5-HT system during birth may lead to impaired 5-HT homeostasis in developing brain regions downstream of DRN and affect their maturation, thereby increasing the risk for disease later on.

## RESULTS

### AVP increases the excitatory synaptic drive onto neonatal 5-HT neurons via V_1A_ receptors

In adult mouse DRN, AVP has been shown to excite 5-HT neurons through a V_1a_ receptor mediated increase in glutamatergic synaptic input (Rood and Beck, 2014). To determine the effect of AVP on developing DRN 5-HT neurons, we performed whole cell voltage clamp recordings in acute brain slices containing DRN of P10-12 neonatal rats. Recorded neurons were filled with biocytin and slices were processed for immunohistochemistry against tryptophan hydroxylase (TPH), the 5-HT synthesizing enzyme, to identify 5-HT neurons. To find the optimal AVP concentration, we tested the effect of 2, 10, 20 and 200 nM AVP on the frequency of spontaneous excitatory postsynaptic current (sEPSC) in the presence of the GABA_A_ receptor antagonist picrotoxin (100 µM) to block fast GABAergic transmission (Suppl. Fig 1). We found that 10 nM AVP application resulted in the strongest increase in sEPSC frequency. Application of 10 nM AVP for 3 min significantly increased sEPSC frequency in 5-HT neurons by 2.77 ± 0.63 fold (Fig. 1 A, D, p=0.021, N=6, Paired Student’s t-test) and the frequency did not return to baseline level following 7 min washout. The effect of 10 nM AVP on sEPSC amplitude was more variable between neurons and hence was on average not significantly increased vs baseline.

**Figure 1.**
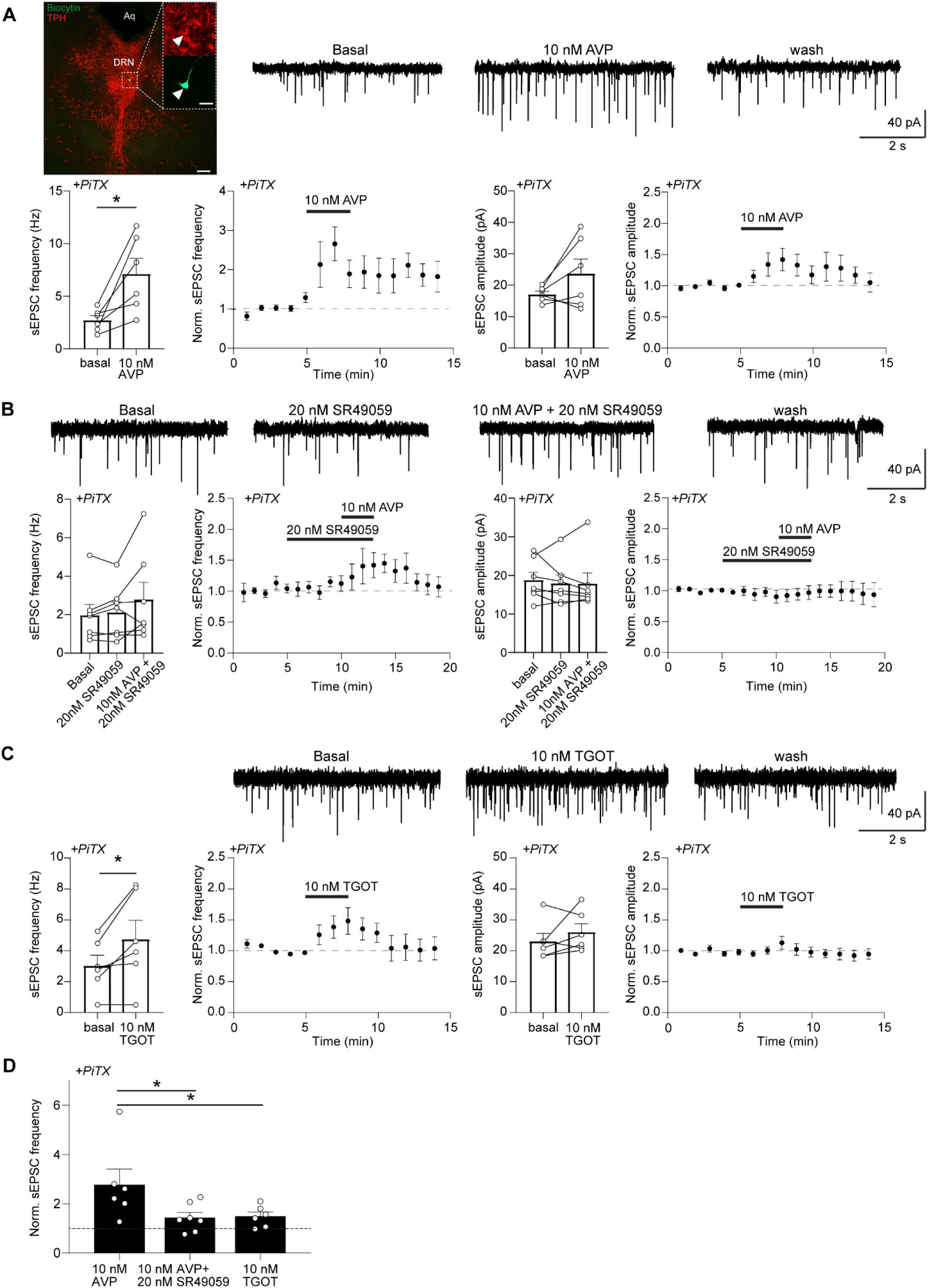
AVP increases sEPSC frequency in neonatal TPH+ neurons in the presence of GABA_A_ antagonist by activating V_1A_ receptors. **A**, Example traces, time-course plots and averaged data (bar diagrams) demonstrating the effect of 10 nM AVP (N=6) on sEPSC frequency and amplitude recorded in the presence of the GABA_A_ antagonist picrotoxin from neonatal TPH+ neurons (N=6) in acute brain slices containing DRN. The representative example image shows a tryptophan hydroxylase (TPH) positive neuron (red, arrowhead) filled with Biocytin (green, arrowhead) in DRN. Aq – aqueduct, scale bars: 200 µm, inset 50 µm. **B**, Same as A for 10 nM AVP + V_1a_ receptor antagonist SR49059 (20 nM, N=7). **C**, Same as A for Oxytocin receptor agonist TGOT (10 nM, N=6). **D**, Bar diagrams comparing the normalised sEPSC frequency (fold increase) between 10 nM AVP, 10 nM AVP + 20 nM SR49059 or 10 nM TGOT. Note the prominent increase in sEPSC frequency by 10 nM AVP that is blocked in the presence of the V_1A_ receptor antagonist. Note also the increase in sEPSC frequency by 10 nM TGOT, but to a much smaller extent than 10 nM AVP suggesting that the effect of 10 nM AVP on sEPSC frequency is mainly mediated by V_1A_ receptors. * *p* < 0.05, Paired Student’s t-test or one-Way ANOVA with post-hoc test.

Next, we tested if this effect was also mainly mediated by V_1a_ receptors as has been described in adult mouse DRN (Rood and Beck, 2014). Application of the selective V_1a_ receptor antagonist SR49059 (20 nM) alone did not affect sEPSC frequency or amplitude (Fig. 1C). Furthermore, combined application of 10 nM AVP in the presence of SR49059 did not significantly affect sEPSC frequency nor amplitude, although a small increase in the sEPSC frequency was observed in few cells (Fig. 1B). Hence, the V_1a_ receptor antagonist blocked the increase in sEPSC frequency by AVP suggesting that the effect is mediated by V_1a_ receptors. Since AVP also binds to oxytocin (OT) receptors (Manning et al. 2012), we tested whether OT receptors are involved in the AVP-mediated increase of sEPSC frequency. Application of the selective OT receptor agonist TGOT (10 nM) led to a small but significant increase of 1.49 ± 0.17 fold in sEPSC frequency in 5-HT neurons (p=0.037, N=6, paired t-test) that returned to baseline during the 7 min washout while sEPSC amplitude was not significantly increased by TGOT (Fig. 1C). However, the magnitude of the increase in sEPSC frequency of 48.84 ± 17.53% was significantly smaller than the increase of 177.30 ± 63.07 % with AVP application (Fig. 1D, One-way ANOVA, p=0.043, with post-hoc test, adjusted p=0.036, N=6-7). This suggests that AVP increases the excitatory synaptic drive onto neonatal 5-HT neurons and that this effect is mainly mediated by V_1a_ receptors with possible minor contribution of the OT receptor.

### AVP increases action potential firing of neonatal 5-HT neurons *in vitro*

To determine, if the AVP-mediated increase in glutamatergic synaptic input onto 5-HT neurons also leads to an increase in their output, we recorded their action potential firing and response to AVP in cell attached mode. Bath application of 10 nM AVP led to a strong and significant 11.96 ± 4.38 –fold increase in action potential firing in all recorded neurons (p<0.001, N=7, Paired student’s t-test) that returned to baseline during the washout period (Fig. 2 A).

**Figure 2.**
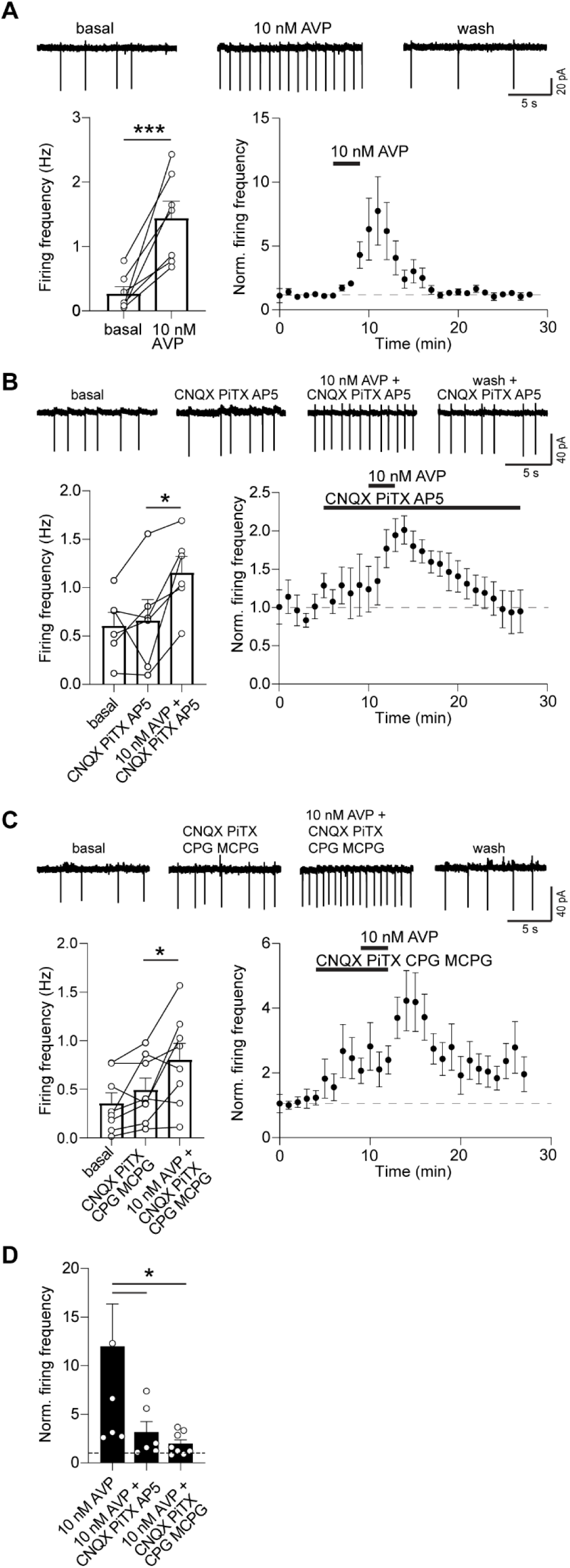
AVP strongly increases spontaneous action potential (AP) firing of neonatal TPH+ neurons in DRN. **A**, Example traces, time-course plots and averaged data (bar diagram) demonstrating the effect of 10 nM AVP (N=7) on spontaneous action potential firing of neonatal TPH+ neurons in acute brain slices containing DRN. **B**, Same as A, for 10 nM AVP in the presence of antagonists blocking fast synaptic transmission (CNQX, Picrotoxin, AP5) (N=6). **C**, Same as A, for 10 nM AVP in the presence of CNQX, Picrotoxin, CPG, MCPG blocking ionotropic and metabotropic glutamate and GABA receptors (N=8). **D**, Bar diagrams comparing the normalised firing frequency (fold increase) under the three conditions between 10 nM AVP, 10 nM AVP + fast synaptic transmission block (CNQX, Picrotoxin, AP5) or 10 nM AVP + ionotropic and metabotropic glutamate and GABA block (CNQX, Picrotoxin, CPG, MCPG). Note the prominent increase in firing frequency by 10 nM AVP that is attenuated but not fully blocked in the presence of glutamate and GABA receptor antagonists, suggesting that at least parts of the effect are mediated by direct actions on 5-HT neurons. * *P* < 0.05, *** *P* < 0.001, Paired Student’s t-test.

To investigate the mechanism underlying the AVP mediated increase in action potential firing, we tested the effect of AVP in the presence of glutamate and GABA receptor blockers (Fig. 2 B, C). Interestingly, co-application of the AMPA/kainate receptor antagonist CNQX, GABA_A_ receptor antagonist picrotoxin and NMDA receptor antagonist AP5 did not affect action potential firing of 5-HT neurons. Additional application of 10 nM AVP led to a 3.14 ± 1.09 fold increase in action potential firing (Fig. 2 B, p=0.023, N=6, paired Student’s t-test) that returned to baseline during the washout period. Similarly, co-application of CNQX, picrotoxin and the selective group I metabotropic glutamate receptor blocker (S)4-CPG and non-selective group I/II (S)-MCPG also did not significantly affect the firing activity of 5-HT neurons (Fig. 2 C). Moreover, additional application of 10 nM AVP was still able to significantly increase action potential firing by 1.95 ± 0.41 fold (p=0.046, N=8, paired Student’s t-test). However, the magnitude of the increase in action potential firing by AVP in the presence of either ionotropic glutamate receptor and GABA receptor blockers or ionotropic and metabotropic glutamate receptor and GABA receptor blockers was significantly smaller than with AVP alone (Fig. 2 D, Kruskal-Wallis test, p=0.026, with post-hoc tests, adjusted p=0.042 or adjusted p=0.011, respectively, N=6-8). This suggests that AVP increases the action potential firing of neonatal 5-HT neurons predominantly through its effects on glutamatergic synaptic transmission onto these neurons. However, the remaining increase in action potential firing by AVP in the presence of those blockers suggests an additional mechanism, comprising most likely a direct effect of AVP on the excitability of neonatal 5-HT neurons.

### AVP does not affect inhibitory synaptic drive onto neonatal 5-HT neurons

To investigate the relative contribution of changes in excitatory versus inhibitory synaptic drive by AVP that underlie the resulting increase in action potential firing, we performed whole cell voltage clamp recordings of glutamatergic sEPSCs and spontaneous GABAergic inhibitory postsynaptic currents (sIPSCs) under experimental conditions where GABAergic events were seen as outward currents and glutamatergic events as inward currents (Fig. 3 A). Under these conditions, 10 nM AVP also significantly increased sEPSC frequency in 5-HT neurons in 5/6 neurons by on average 5.90 ± 1.96 fold (Fig. 3 B, p=0.036, N=6, Paired student’s t-test) while sEPSC amplitude was not increased (Fig. 3 C). The baseline rate of sIPSCs was very low at this age (0.15 ± 0.02 Hz). The effect of AVP on sIPSCs frequency was variable and neither changes in frequency nor amplitude were significant. However, one neuron showed a strong increase in sIPSC frequency in response to 10 nM AVP whereas the remaining five neurons were not responding.

**Figure 3.**
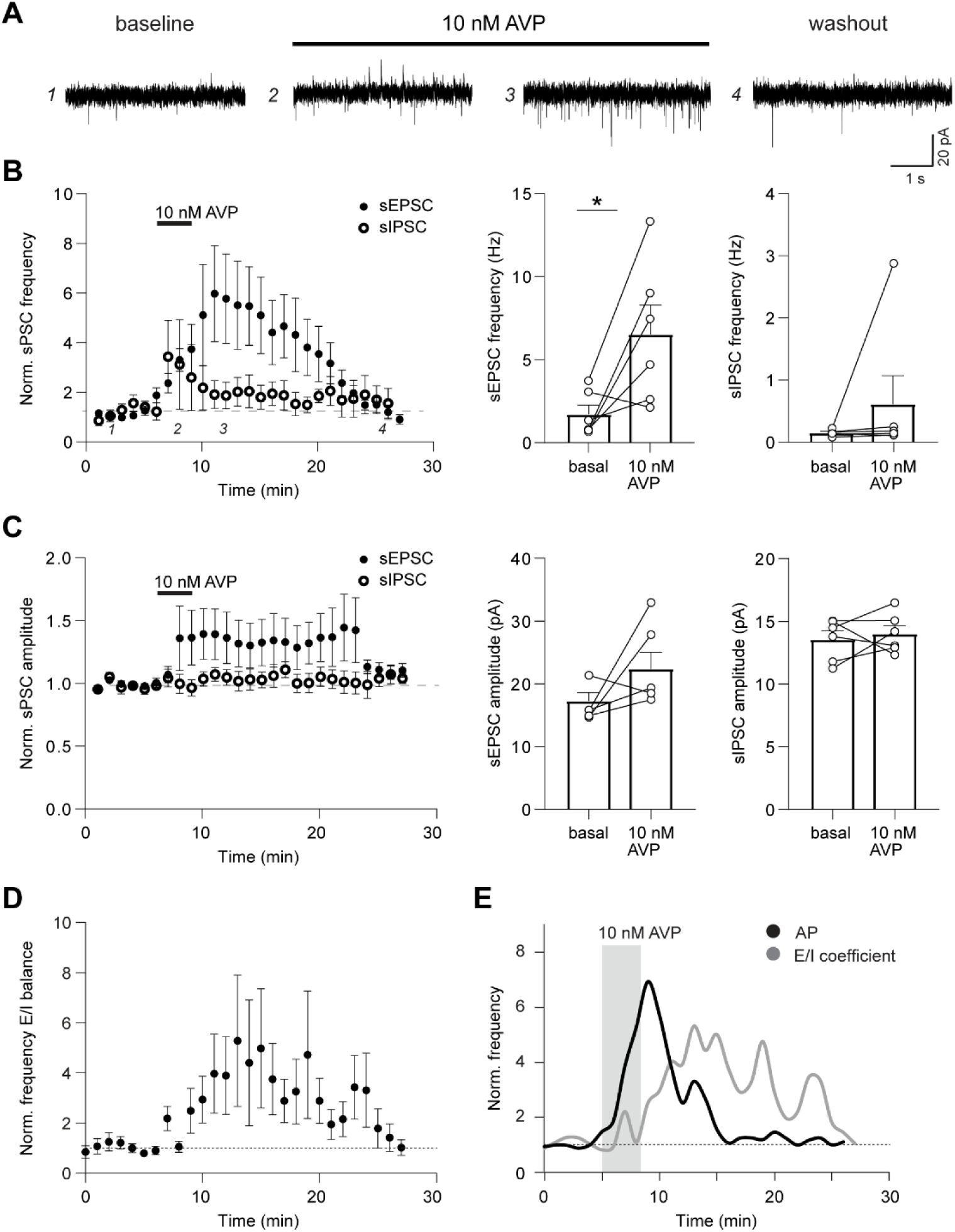
AVP does not affect GABAergic synaptic transmission onto neonatal 5-HT neurons. **A**, Representative example traces illustrating effect of 10 nM AVP on the frequency and amplitude of sEPSC (inward) and sIPSC (outward), recorded simultaneously at holding potential –47 mV from neonatal TPH+ neurons in acute brain slices containing DRN. Numbers mark baseline (1), peak in sIPSC in response to AVP (2), peak in sEPSC in response to AVP (3), and washout (4), respectively. **B**, Time-course plot and averaged data demonstrating the effect of 10 nM AVP on sEPSC and sIPSC frequency recorded simultaneously at holding potential –47 mV from TPH+ neurons (N=8) in acute brain slices containing DRN. Quantification of the effect of AVP on the frequency of sEPSCs (bar diagrams) corresponds to time point 3, whereas quantification of the effect of AVP on of the frequency of sIPSCs (bar diagrams) corresponds to time point 2. **C**, Same as B for sEPSC and sIPSC amplitude. **D**, Time-course plot demonstrating effect of 10 nM AVP on E/I balance of PSC frequency in TPH+ neurons (N=8) calculated as the ratio of sEPSC/sIPSC frequency in acute brain slices containing DRN. **E**, Summary schematic illustrating effect of 10 nM AVP (grey bar) over time on normalized spontaneous AP firing frequency (black line, N=8) and E/I coefficient (grey line, N=8) in TPH+ neurons in in acute brain slices containing DRN. Note that the effect on AP firing frequency is preceding the effect on E/I coefficient, suggesting that the increase in AP firing is mainly mediated by direct actions of AVP on TPH+ neurons. * *P* < 0.05, Paired Student’s t-test.

Interestingly, when plotting the time course of the response of action potential firing (see Fig. 2A) and E/I coefficient to AVP application (Fig. 3 D), the increase in AP firing frequency clearly precedes the peak in E/I coefficient (Fig. 3 E). This further supports the notion that in addition to the effects on glutamatergic transmission, the effect of AVP on action potential firing is likely to involve also direct actions on neonatal 5-HT neurons.

### Neonatal DRN 5-HT neurons show typically tonic regular or low frequency oscillation firing pattern *in vivo*

In the adult rodent brain, 5-HT neurons in DRN are characterized by a slow tonic regular or clock-like firing pattern under urethane anaesthesia (Allers and Sharp, 2003). Furthermore, bursting 5-HT neurons that exhibit regular firing activity but fire doublets or triplets of action potentials have also been described (Hajos et la., 2007; Schweimer and Ungless, 2010; Schweimer et al., 2011). To investigate whether developing 5-HT neurons display similar firing activity, we performed *in vivo* electrophysiological single unit recordings combined with juxtacellular labelling in P10-12 urethane anaesthetised rat pups. A total of 39 DRN neurons in 28 animals were successfully Neurobiotin (NB) labelled and processed for immunohistochemistry against 5-HT. 25/39 neurons were immunopositive for 5-HT (5-HT+). These 5-HT+ neurons had predominantly two firing patterns: 9/25 (36%) had tonic clock-like firing patterns (Fig. 4 A) similar to the typical 5-HT neurons in the adult rodent brain. Their firing rates were low (median rate 0.73 (0.65) Hz) but very regular (mean coefficient of variation (CoV) of the inter-spike interval (ISI) 0.35 ± 0.02). Accordingly, they showed clear peaks in the autocorrelation histogram (Fig. 4 Aiii) as well as a narrow distribution of ISIs (Fig. 4 Aiv) which is typical for regular firing neurons. They also displayed a broad spike waveform (3.45 ± 0.14 ms) which is also characteristic of adult 5-HT neurons. Interestingly, neonatal tonic-firing 5-HT neurons also exhibited pauses in firing activity of 49 ± 6.5 s on average.

**Figure 4.**
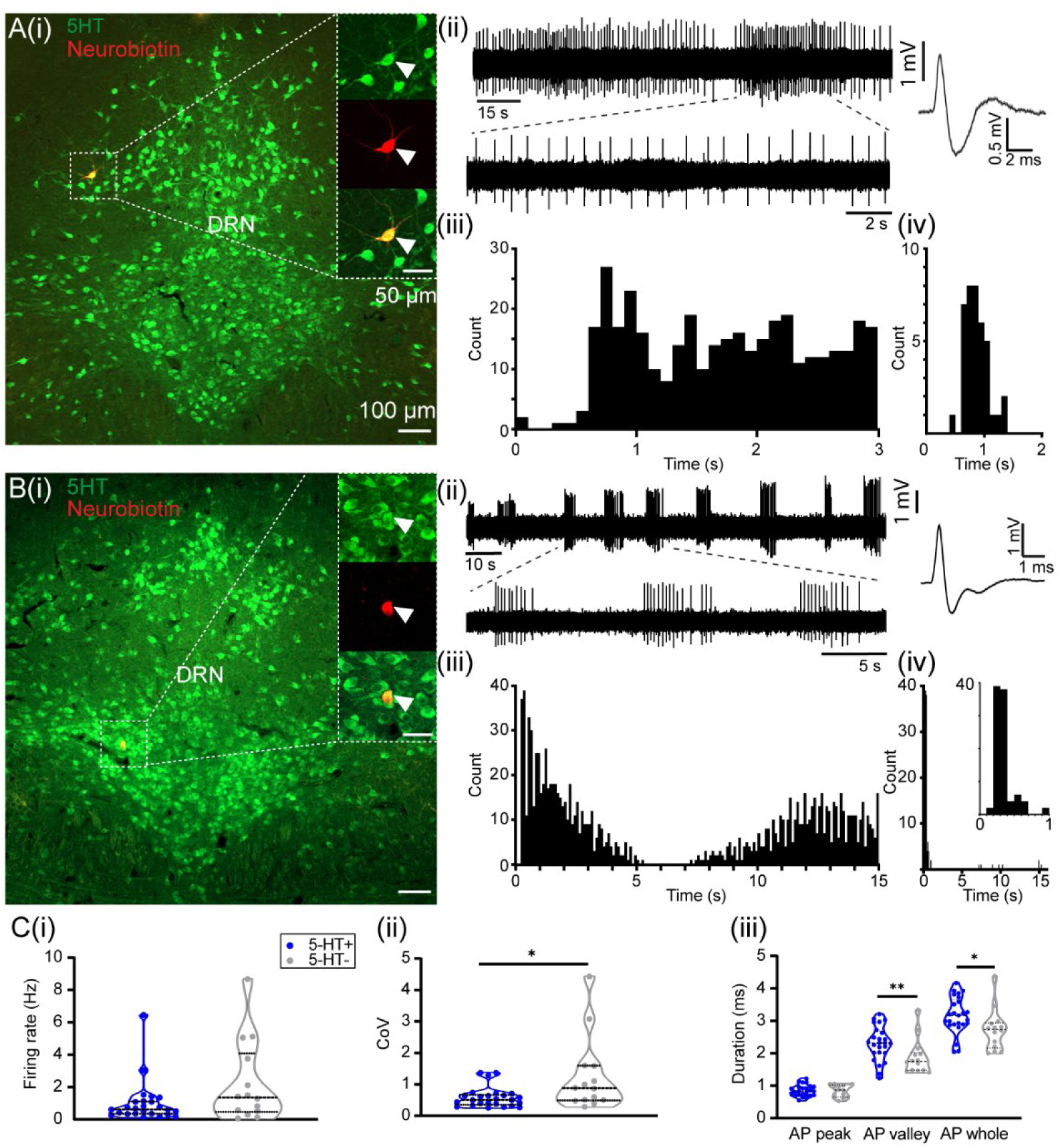
Tonic regular and slow oscillating activity are typical firing patterns of neonatal 5-HT neurons *in vivo*. **A**, Tonic regular firing neuron. Photomicrograph **(i)** of 5-HT-positive neuron in neonatal dorsal raphe nucleus (DRN) (green, arrowhead) labelled with Neurobiotin (red, arrowhead) during *in vivo* electrophysiology. **(ii)** Corresponding spike train and waveform, **(iii)** autocorrelation histogram of spikes (bin size 100 ms) and **(iv)** inter-spike interval (ISI) histogram (bin size 100 ms) of the labelled neuron in **(i)**. Note the wide action potential (AP) and tonic regular firing pattern (see enlarged trace) as characterized by a peak in the autocorrelation (0.6-1.1 s) and narrow distribution of ISIs typical for 5-HT neurons **(iv)**. **B, Neuron displaying low-frequency oscillations (LFO) in firing.** Same as **A (i-iv)** for LFO neuron. Note the typical wide AP waveform and oscillating occurrence of regular spike trains at low frequency (see enlarged trace) as characterised by peaks in the autocorrelation (bin size 100 ms) and ISI histogram (bin size 100 ms) at both short (see inset in ISI histogram) and longer timescales. **C,** Violin plots of **(i)** firing rate, **(ii)** CoV and **(iii)** and action potential properties of 5-HT+ (blue dots, n=25) and 5-HT-(grey dots, n=14) neurons. Note the significantly lower CoV of 5-HT+ neurons indicating more regular firing activity as well as longer AP valley indicating a longer afterhyperpolarization resulting in longer duration of the whole AP compared to 5-HT-neurons. * *P* < 0.05, ** *P* < 0.01, Unpaired t-test or Wilcoxon rank sum test.

The largest group of neonatal 5-HT+ neurons, 11/25 (44%), were characterized by low frequency oscillations (LFOs) in firing rate. These neurons fired short trains of action potentials that occurred at regular or irregular intervals. The firing pattern within the train was regular (CoV 0.61 ± 0.05) at low rate (median 0.84 (2.06) Hz) and spike trains occurred at an average interval of 16.21 ± 3.42 s (range 3.00 –38.89 s). Similar to the tonic regular firing 5-HT+ neurons, they also had a broad action potential waveform (2.94 ± 0.20 ms). Figure 4 B shows an example of a neuron that fires regular trains every 10-15 s. The autocorrelation histogram displays peaks at both shorter time scales (see peaks at 0.2-0.4s, 0.5-0.7s and 0.8-1s in Fig. 4 BiV) corresponding to spikes fired within the train and broad peaks at longer time scales corresponding to the regular occurrence of the train e.g. every 10-15 s in this example. Correspondingly, the ISI histogram also shows a narrow peak at short time scales (see inset Fig. 4 BiV) as well as larger ISIs above 7 s that correspond to the regular pauses in firing. In addition, three 5-HT+ neurons (12%) that changed their firing activity from LFOs to more tonic firing during the recording period were also observed. We also recorded a single bursting 5-HT+ neuron (4%) that fired spike doublets at tonic regular rate (Suppl. Fig. 2 A) and a single 5-HT+ irregular neuron (4%) that fired at very low rate.

A further 14/39 neurons were immunonegative for 5-HT (5-HT-) that displayed a wide range of firing patterns. Most common were irregular firing neurons (6/14, 42.9 %) that fired at low rate <1 Hz. A further 3/14 (21.4 %) neurons were fast firing neurons with a mean rate >5 Hz, three neurons (21.4 %) showed tonic regular firing activity and two neurons (14.3 %) showed LFOs in their firing activity. When comparing the firing properties of 5-HT+ vs 5-HT-DRN neurons (Fig. 4 C), there was no significant difference in their firing rate (0.61 (0.76) Hz vs 1.35 (3.61) Hz). However, 5-HT+ neurons had a significantly lower CoV than 5-HT-neurons (0.51 (0.32) vs 0.88 (1.10), p=0.0262, N=14-25, Wilcoxon rank sum test,), i.e. their firing activity was more regular. When comparing their action potential (AP) waveforms, 5-HT+ neurons had a significantly broader AP than 5-HT-negative neurons (3.17 ± 0.11 vs 2.72 ± 0.17 ms, p=0.0274, N=14-25, unpaired student’s t-test) due to significantly longer afterhyperpolarization (AP valley, 2.32 (0.66) ms vs 1.75 (0.54) ms, p=0.0098, N=14-25, Wilcoxon rank sum test), while the initial AP peak was not different between the groups.

Hence, neonatal 5-HT neurons in DRN display predominantly two main firing patterns, slow tonic regular firing or LFOs of regular spike trains. They can be distinguished from non-5-HT neurons by their broad AP waveform and particularly longer afterhyperpolarization as well as by their more regular firing activity.

### Neonatal putative 5-HT neurons in DRN are activated by AVP *in vivo*

To confirm our *in vitro* findings and test the response of neonatal DRN 5-HT neurons to AVP *in vivo*, we carried out multi-channel *in vivo* local field-potential and multi-unit recordings in DRN combined with local drug or vehicle injections in urethane-anaesthetised neonatal rats (Fig. 5 A). We recorded baseline neuronal activity in DRN for 15 min before injection of 200 nl of 0.9% NaCl (vehicle control) or 100 nM AVP, followed by a further 15 min of recording after the injection. After pre-processing the data and sorting the recorded spikes into single units, we had n=49 stable units from 10 rats in the vehicle control group and n=95 units from 10 rats in the AVP group. In the vehicle control group, 6 neurons (12%) were excited and increased their firing rate, 13 (26%) neurons were inhibited and decreased their firing rate and an additional 30 units (62%) showed no change in firing rate i.e. were not responding to vehicle injection (Fig. S3). This ratio of excited, inhibited and non-responding neurons was significantly different in the AVP group (Chi-square test, p<0.0001), where 36 neurons (38%) were excited, 35 neurons (36 %) were inhibited and an additional 25 neurons (26 %) were unresponsive to the drug (Fig. 5 B). Hence, in the vehicle control group, the majority of neurons did not respond to vehicle injection, while some neurons were inhibited and very few were excited. In contrast, local injection of AVP led to an excitatory response in the largest fraction of recorded neurons while less neurons were inhibited or did not respond. Baseline firing rates of excited (median 1.23 (1.72) Hz), inhibited (2.67 (3.74) Hz) and non-responding neurons (2.28 (2.85) Hz) were low.

**Figure 5.**
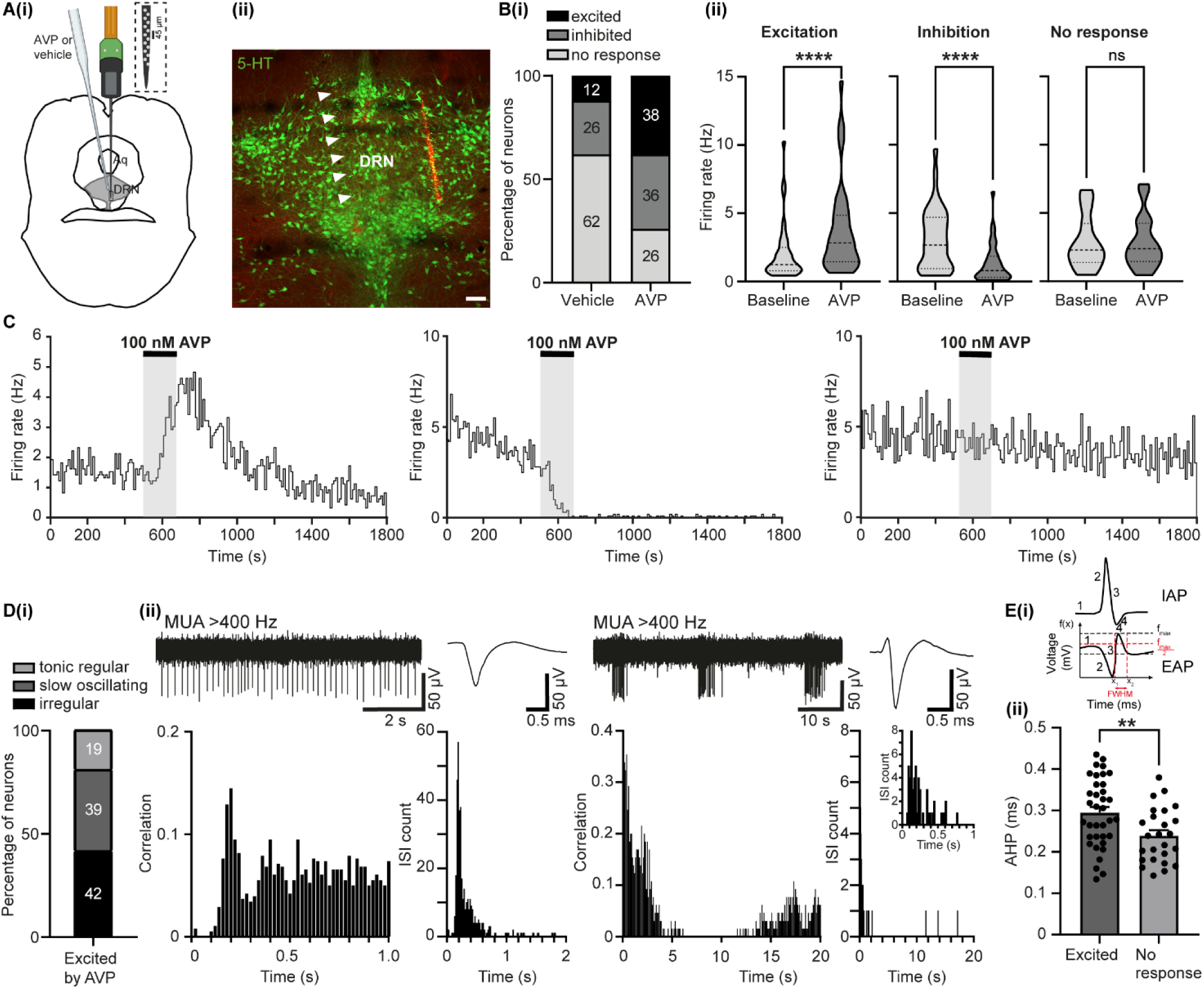
AVP increases firing activity of putative 5-HT neurons in DRN *in vivo.* **A**, (i) Diagram illustrating the 64-channel recording of neuronal activity in DRN combined with local vehicle or drug injection via an angled glass pipette. Inset shows polytrode channel layout with 45 µm distance between individual channels. **(ii)**, Photomicrograph image of coronal rat brain section showing the labelled tract of the recording electrode (red) and unlabelled tract of the injection cannula (arrowheads) within DRN. Immunohistochemical staining against 5-HT (green) shows location and boundaries of DRN. Scale bar: 100 µm. **B, (i)** Bar diagram showing the percentage of excited, inhibited and not responding neurons following local vehicle or AVP injection in DRN. The distributions within each treatment group were significantly different (Chi square test, P<0.0001). **(ii)** Violin plots of firing rates during baseline and after 100 nM AVP injection for excited (N=36, left), inhibited (N=34, center) and not responding (N=25, right) DRN neurons. **C**, Firing rate histograms over time (bin size= 10 s) of individual representative DRN neurons responding with increase (left) or decrease (center) or showing no response (right) to local injection of 100 nM AVP (grey shaded area). **D, (i)** Bar diagram showing the distribution of tonic regular, LFO and irregular firing DRN neurons excited by AVP. **(ii)** Example trace, average waveform, autocorrelation histogram and ISI histogram (bin size= 20 ms) of a tonic regular firing neuron. Note the clear peak in the autocorrelation at lag 200 ms as well as the narrow distribution of ISIs typical for a regular firing neuron. **(iii)** Same as (ii) for a LFO neuron. Note the bimodal distributions of autocorrelation histogram (bin size 100 ms) at short and long lags as well as ISIs (bin size 20 ms) corresponding to the regular firing of spikes within a train (see inset at enlarged time scale) and the slow regular occurrence of these spike trains every 15-20 s in this example. **E, (i)** Diagram of intracellular action potential (IAP) and its 1^st^ derivative, the extracellular action potential (EAP) of opposite polarity below with the typical phases of 1) resting membrane potential, 2) depolarisation, 3) repolarisation and 4) hyperpolarisation. To analyse the duration of the afterhyperpolarisation phase (AHP) as shown in the bar diagram below, we calculated the full width at half maximum (FWHM) which corresponds to the width of a spectral peak at half of its maximum amplitude. This can be visualized as the distance between two points on the x-axis (x_1_ and x_2_) where the curve’s intensity is half of its peak value. **(ii)** Bar diagram showing the AHP of DRN neurons excited by AVP *in vivo* or showing no response. Note that excited neurons have significantly longer AHP than non-responding neurons. **** p<0.0001, **p<0.01, Wilcoxon matched-pairs signed rank test or unpaired Student’s t-test.

Interestingly, baseline firing rates of AVP-excited neurons were significantly lower than AVP-inhibited or AVP-unresponsive neurons (Kruskal-Wallis test, p=0.025, multiple comparisons tests, adjusted p=0.016 and p=0.016, respectively). Excited neurons increased their firing rates by 1.59 (0.96) fold (p<0.0001, N=36, Wilcoxon matched-pairs signed rank test) whereas inhibited neurons decreased their firing rates by 0.47 (0.46) fold in response to AVP (Fig. 5 B, (p<0.0001, N=34, Wilcoxon matched-pairs signed rank test, Fig. S3). Figure 5C shows representative examples of the change in firing rate of individual neurons in response to AVP. In comparison to the vehicle injections that did not affect the firing rate of the majority of neurons tested, the predominant response of DRN neurons to AVP is excitation. Whether AVP also leads to inhibition of DRN neurons *in vivo* is less certain, as local injection of a vehicle also reduced the firing rate of some of the recorded neurons. Since our *in vitro* electrophysiological recordings showed a clear excitation of identified 5-HT neurons in response to AVP, we analysed if DRN neurons excited by AVP *in vivo* also share characteristics typical of the above identified neonatal 5-HT neurons *in vivo* (see Fig. 4). Indeed, we found that of the 36 neurons excited by AVP, seven neurons (19%) were tonic regular firing neurons, 14 neurons (39%) showed LFOs in firing activity and 15 neurons (42%) had an irregular firing pattern (Fig. 5 D). As shown in Fig. 4 above, tonic regular firing and LFOs are the predominant firing patterns of identified neonatal 5-HT neurons *in vivo*. Tonic regular firing DRN neurons excited by AVP fired at low rate (1.9 (1.97) Hz) and also displayed a similar peak in the autocorrelation histogram, the narrow ISI distribution and a broad action potential waveform typical of identified 5-HT neurons *in vivo* (Fig. 5Dii). Neurons with LFO firing activity excited by AVP fired regular trains of action potentials at regular intervals of 20-40 s (baseline rate 1.55 (4.06) Hz) similar to those identified as 5-HT neurons by juxtacellular labelling (see Fig. 4). These neurons also show a bimodal distribution in the autocorrelation or ISI histograms corresponding to regular firing within a spike train as well as an oscillatory occurrence of these trains at low frequency over time. Finally, when comparing the duration of the afterhyperpolarization phase (AHP) between AVP-excited or not responding neurons, excited neurons had significantly longer AHP than non-responsive ones (Fig. 5 E, 0.29 ± 0.01 ms vs 0.24 ± 0.01 ms, p=0.0069, unpaired t-test, N=25-36). The long afterhyperpolarization of the AP waveform is also a characteristic feature of identified neonatal 5-HT neurons *in vivo* (see Fig. 4).

Our results show that the main effect of AVP on the firing activity of DRN neurons *in vivo* is excitatory and that a large proportion (up to 60%) of the excited cells have the typical firing patterns and broad AP waveform of previously identified neonatal 5-HT neurons.

### The neonatal DRN has a sparse vasopressinergic innervation with strong sex dependence

The adult mouse DRN is characterised by a moderate-to-dense network of vasopressinergic fibers and terminals most prominent in the dorsal subregion (DeVries et al., 1985; Rood and De Vries, 2011; Rood and Beck, 2014). The innervation is sexually dimorphic with males having a denser fiber plexus than females (Rood et al., 2013). However, to our knowledge, no studies have been conducted to characterize the vasopressinergic innervation of DRN during neonatal development.

We performed immunohistochemistry against AVP-associated neurophysin2 on coronal brain sections containing DRN of P10-12 neonatal male and female rats to label and analyse the neonatal vasopressinergic innervation of DRN. In males, we typically observed a still sparse innervation of pronounced thick fibers as well as a loose network of thin fibers with boutons (Fig. 6 A, B). When quantifying the fractional areas covered by AVP immunoreactive fibers throughout DRN subregions, we found similar fiber fractional area percentages (FFAPs) in rostral (0.29 ± 0.02, n=39 images from N=7 pups), central (0.27 ± 0.02, n=56 images from N=7 pups) and caudal DRN (0.24 ± 0.03, n=27 images from N=7 pups). However, the previously described predominant vasopressinergic innervation in dorsal DRN was already present at neonatal age with significantly higher FFAP in dorsal (0.29 ± 0.02, n=54 images from N=7 pups) versus ventral DRN (0.19 ± 0.02, n=40 images from N=7 pups, p=0.0001, unpaired t-test). In contrast, the female neonatal DRN was almost devoid of AVP-ir fibers at this age with often only a single fiber visible. Accordingly, when comparing the pooled FFAP of AVP-ir fibers for whole DRN between males and females, males had significantly higher FFAP than females (Fig. 6 C, 0.27 ± 0.02 vs 0.05 ± 0.01, N=7/sex, p<0.0001, unpaired t-test).

**Figure 6.**
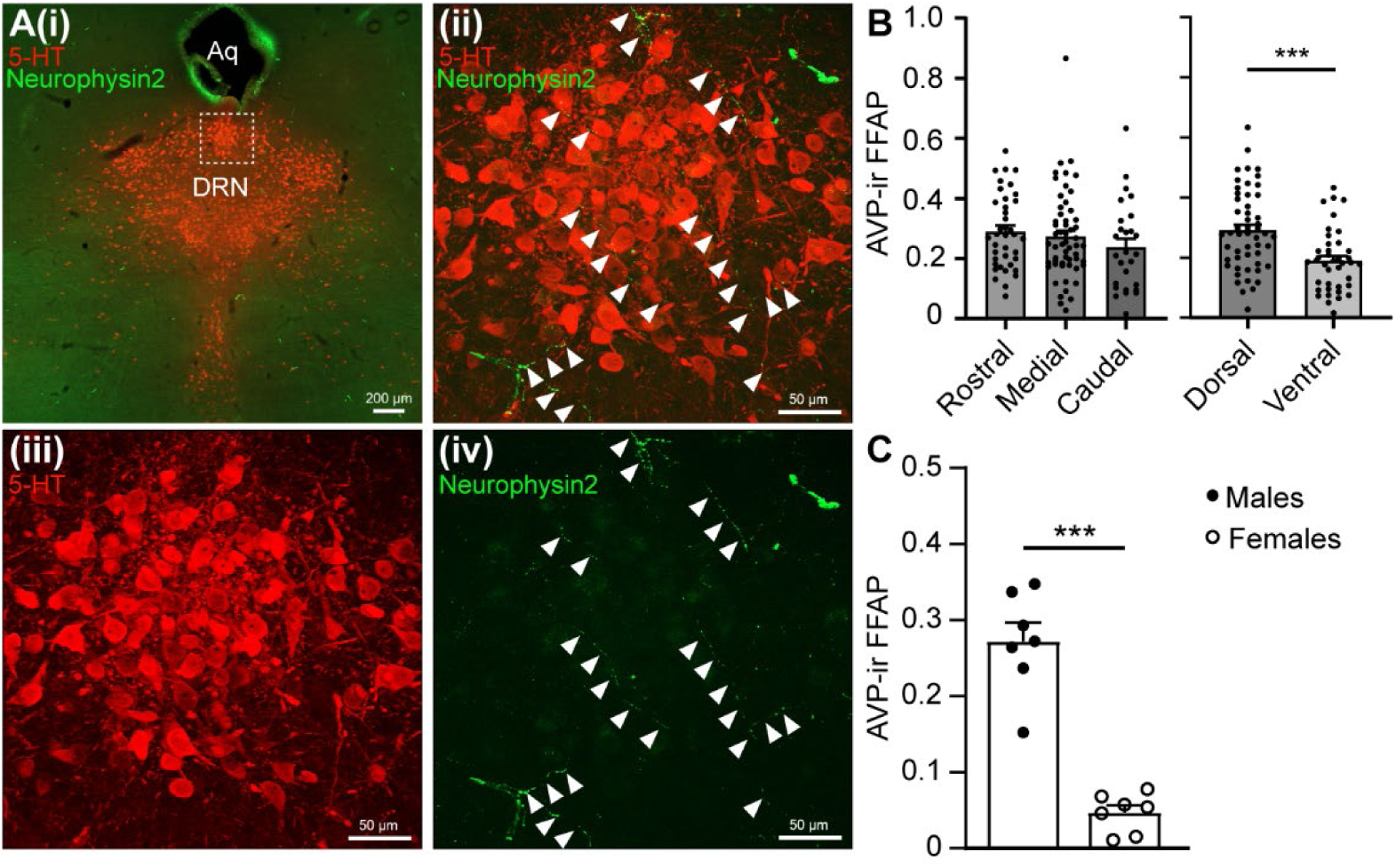
The vasopressinergic innervation of the rat DRN is already present during neonatal development in males. **A**, (**i**) Photomicrograph of a coronal brain section from a male P11 rat pup including the DRN with immunostaining against 5-HT (red), determining the outline of DRN and immunolabelling against neurophysin2 (green). (**ii**), Double immunolabelling against 5-HT (red) and neurophysin2 (green) of same area in dorsal DRN as marked by white box in (i). Note the presence of neurophysin2 positive fibers (arrowheads) in the vicinity of 5-HT neurons. (**iii**) Same image as in (ii) showing only 5-HT staining. (iv) Same image as in (ii) showing only neurophysin2-positive fibers (arrowheads). Scale bar in (i) =200 µm, in (ii-iv) =50 µm. **B**, Bar diagrams showing FFAP of neurophysin2 for whole DRN in male (n=7) versus female (n=7) rat pups. Note the significantly larger FFAP in males. **C**, Bar diagrams showing FFAP of neurophysin2 in male pups (n=7) for rostral (n=39 images, 5-6 per section/pup), medial (n=56 images, 8 per section/pup) and caudal (n= 27 images, 3-4 per section/pup) DRN (left) as well as for dorsal (n=54 images) vs ventral (n=40 images) DRN. Note that there is no difference in FFAP in the rostral to caudal dimension but a higher FFAP in dorsal DRN corresponding to a more dense innervation in dorsal compared to ventral DRN. *** *p* < 0.0001, unpaired t test.

This suggests that the density of vasopressinergic fibers in DRN is still sparse during neonatal age and undergoes developmental change towards moderate to dense innervation at adulthood. However, the predominant presence of AVP-ir fibers in dorsal DRN as well as the strong sex dependence of the vasopressinergic innervation of DRN are already present at neonatal age.

### The vasopressinergic innervation of the neonatal DRN in the rat originates exclusively from cell groups in medial amygdala and BNST

In the rodent brain, AVP neurons are located in five brain regions: the paraventricular nucleus of the hypothalamus (PVN), suprachiasmatic nucleus (SCN), supraoptic nucleus (SON), bed nucleus of the stria terminalis (BNST) and medial amygdala (MeAmy). In BNST and MeAmy AVP production is dependent on gonadal steroids and gonadectomy in both male and female rodents leads to a depletion of AVP in their projection sites (de Vries et al., 1984; Mayes et al., 1988; Miller et al., 1992; Wang and De Vries, 1995). Using this indirect approach, the vasopressinergic innervation of DRN has been proposed to originate mainly from BNST and MeAmy, since gonadectomy led to a decrease in AVP-immunoreactivity in the DRN in adult male and female mice (Rood et al., 2013). Very recent elegant viral tracing experiments confirmed that both AVP neurons in BNST and MeAmy project to DRN in adult mice (Rigney et al., 2023). However, it remained unknown whether 1) this is also true in rats, 2) whether the innervation is exclusive from BNST and MeAmy and 3) whether the fiber origin is the same during early postnatal development.

To address these issues, we performed classical retrograde tract tracing experiments combined with immunohistochemistry against AVP-associated neurophysin-2. We performed iontophoresis of the retrograde tracer Fluorogold (FG) into DRN of P12 male rat pups. After four days survival time, animals were sacrificed by perfusion-fixation, brains extracted and sectioned for further immunohistochemistry. In total, N=5 experiments were included in the analysis with correct location of the FG injection site within the boundaries of DRN (Fig. S4). We imaged and analysed all sections containing neurophysin-2-ir or FG-ir cells in MeAmy, BNST, PVN and SON (Fig. 7, Table 2). In MeAmy and BNST (see Fig. 7 A), we found scarce cell groups of neurophysin-2 positive neurons in posterodorsal MeAmy and in medial posteromedial (BSTMPM) and medial posterolateral BNST (BSTMPL) as previously reported (Rood and De Vries, 2011) that were very similar in total number in both regions (Table 2).

**Figure 7.**
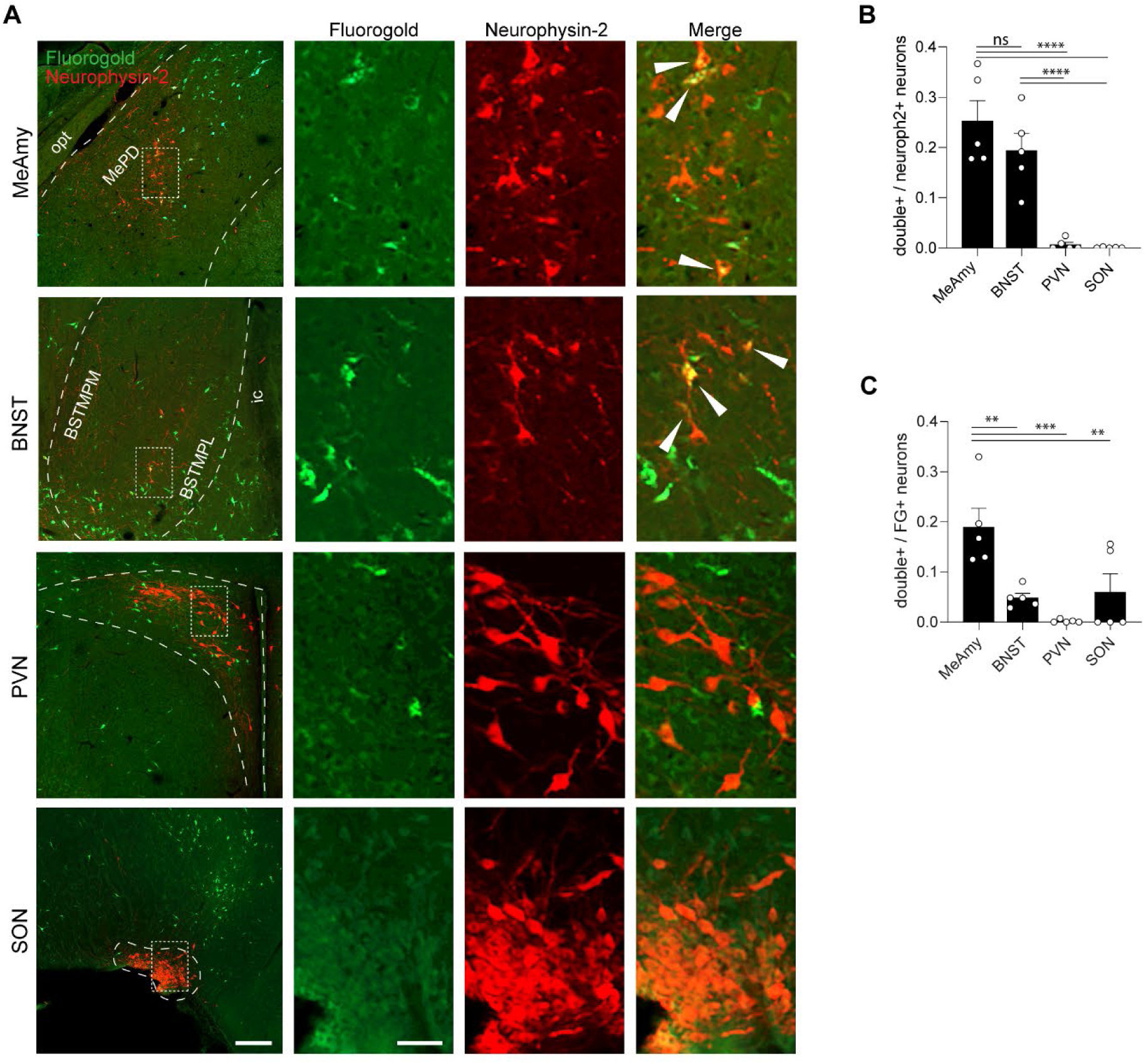
The vasopressinergic innervation of the neonatal dorsal raphe nucleus originates exclusively from cell groups in medial amygdala and BNST. **A**, Example images and insets illustrating Fluorogold positive (FG+, green), Neurophysin-2 positive (Neuroph-2+, red) and double positive neurons (merge, yellow) in medial amygdala (MeAmy) posterodorsal part (MePD), bed nucleus of stria terminalis (BNST) medial posteromedial (BSTMPM) and medial posterolateral (BSTMPL) part, paraventricular nucleus (PVN) and supraoptic nucleus (SON). BNST, MeAmy, PVN and SON are outlined. Insets are marked with a rectangle. FG and Neuroph-2 double labelled neurons are marked with an arrowhead on merge inset. Scale bars 200 µm, 50 µm on insets. **B**, Bar diagram depicting the ratio of Neuroph2+ neurons that are FG and Neuroph2 double+ analysed in MeAmy, BNST, PVN or SON, .ie. the ratio of Neurophy2+ neurons that project to DRN in MeAmygd, BNST, PVN or SON. N (animals) = 5. **C**, Bar diagram depicting the ratio of FG+ neurons that are FG and Neuroph2 double+ analysed in MeAmy, BNST, PVN or SON, .ie. the ratio of FG+ neurons that project to DRN and also contain AVP in MeAmygd, BNST, PVN or SON. N (animals) = 5. * *p* < 0.05, ** *p* < 0.01, *** *p*< 0.001, **** *p*<0.0001, One-way ANOVA with FDR-corrected multiple comparison tests, opt: optic tract, ic: internal capsule.

In line with data from adult mice, we found double FG/neurophysin-2 immunopositive neurons (double+ neurons) nearly exclusively in BNST and MeAmy (Fig. 7 B). On average 25.27 ± 4.05 % (range 17.75-36.66 %) or 19.39 ± 3.47 % (range 9.07-29.91 %) of neurophysin-2+ neurons were also positive for FG, i.e. projected to DRN in MeAmy or BNST, respectively and these ratios were significantly larger than those for PVN or SON (One way ANOVA, p<0.0001, multiple comparison tests, adjusted p<0.0001). However, these ratios were not significantly different between MeAmy and BNST suggesting that the vasopressinergic innervation at least in neonatal male rats originates from both MeAmy and BNST with approximate equal contribution. In PVN or SON, we only found double positive neurons in 2/3 animals, and only very low numbers i.e. one animal in each region with a single double positive neuron and one animal each with 5 or 6 double positive neurons. Accordingly, on average 0.67 ± 0.48 % neurons in PVN or 0.07 ± 0.05 % neurons in SON were double positive suggesting that the contribution of AVP neurons in PVN and SON to vasopressinergic innervation in DRN is negligible.

However, MeAmy, BNST and PVN had on average many FG+ neurons with highest numbers in BNST (Table 2) showing that the vasopressinergic innervation is only a fraction of other afferent input from these regions to DRN. On average 18.96 ± 3.73 % of FG+ neurons were also positive for neurophysin-1 in MeAmy, 4.85 ± 0.90 % in BNST, 0.19 ± 0.14 % in PVN and 5.97 ± 3.66 % in SON (Fig. 7 C). Accordingly, of those neurons projecting to DRN, MeAmy contained the highest proportion of vasopressinergic cells that was significantly larger than in all other analysed regions (One way ANOVA, p=0.0009, multiple comparison tests adjusted p=0.0051, p=0.0008, p=0.0064). However, most of the innervation to DRN from MeAmy and BNST is from other neurons than those containing AVP, while in PVN the innervation of DRN seems to exclusively originate from non-vasopressinergic neurons. SON does not seem to project to DRN at all based on the rather low numbers of FG+ cells.

These results show that the vasopressinergic innervation of the DRN in neonatal rats originates in MeAmy and BNST with a very similar contribution of either region. Neither vasopressinergic neurons in PVN nor SON seem to innervate DRN. These results in rats are in line with previous studies using gonadectomy (Rood et al., 2013) or viral tracing (Rigney et al., 2023) in adult mice to determine AVP fiber origin in DRN and show that MeAmy and BNST are the main source of AVP also in the neonatal rat DRN.

## Discussion

The present study indicates that the inhibitory function of AVP, previously characterized in cortical networks at birth, is not universal in the brain and that AVP can also be strongly excitatory in subcortical neonatal networks such as the DRN 5-HT system. Our data show that AVP increased the excitatory synaptic drive in neonatal 5-HT neurons via V_1A_ receptors while inhibitory synaptic input remained largely unaffected. Accordingly, AVP also prominently increased action potential firing of neonatal 5-HT neurons by enhanced glutamatergic synaptic input and most likely also by a direct facilitation of the excitability of 5-HT neurons. Identified neonatal 5-HT neurons showed two predominant firing patterns *in vivo*, either slow tonic regular firing similar to the firing activity described in adulthood or low frequency oscillations of short regular spike trains. The majority of putative neonatal 5-HT neurons *in vivo* also responded with an increase in firing activity to local AVP injections. The vasopressinergic innervation in neonatal DRN was still sparse and originated exclusively from cell groups in BNST and medial amygdala. Therefore, while a fine balance between inhibitory function of AVP in cortical versus excitation in subcortical networks may protect the brain at birth, exaggerated AVP release during birth stress may cause this balance to break down. This could lead to strong activation of the neonatal 5-HT system affecting its own ongoing functional development as well as 5-HT homeostasis in target regions that may contribute to an increased risk to disease later on in life.

### AVP is strongly excitatory in the neonatal dorsal raphe 5-HT system

Our data clearly show that AVP has a powerful excitatory effect on 5-HT neurons in the neonatal rat DRN by increasing glutamatergic synaptic input in a predominantly V_1a_ receptor dependent manner. This is in line with data from adult mouse DRN, where AVP has been shown to increase both frequency and amplitude of EPSCs in 5-HT neurons via V_1a_ receptors (Rood and Beck, 2014) while OT receptors were not involved. In our experiments we cannot fully exclude the contribution of OT receptors, since application of a selective OT agonist also led to a slight but significant increase in sEPSC frequency but much smaller in magnitude than by AVP. Since pharmacologically blocking V_1a_ receptors prevented the increase in sEPSC frequency by AVP, the effect seems to be mainly mediated by these receptors while OT receptors may contribute to the effect.

AVP also strongly increased action potential (AP) firing in neonatal DRN 5-HT neurons. Interestingly, application of ionotropic and metabotropic glutamate receptor blockers and GABA_A_ receptor blockers did not affect the firing frequency of neonatal 5-HT neurons *in vitro.* This suggests that in this immature stage neither glutamatergic nor GABAergic synaptic inputs are required for maintaining spontaneous AP firing and that this governed through other mechanisms which may involve noradrenergic regulation of firing by α1 receptors beyond others (for review see Maejima et al., 2013). Furthermore, application of these blockers did neither prevent an increase in the firing frequency of 5-HT neurons by AVP. However, the magnitude of the increase in the presence of the blockers was significantly smaller than with AVP alone. Accordingly, AVP may boost AP firing by a combination of increasing glutamatergic synaptic inputs and a direct enhancement of the excitability of neonatal 5-HT neurons. Rood and Beck, 2014 previously suggested that the excitatory effect of AVP on adult mouse 5-HT neurons is indirect and relayed via glutamatergic neurons within the DRN. Recent data on V_1a_ receptor expression in adult mouse DRN showed that at least three populations of neurons express these receptors, glutamatergic, glutamatergic/5-HT and GABAergic neurons (Patel et al., 2022). Hence it is likely that the same populations of neurons also express these receptors in neonatal DRN and that AVP acts on local glutamatergic neurons to increase excitatory synaptic input of 5-HT neurons and drive AP firing. In addition, AVP is also known to increase the excitability of neurons directly via V_1a_ receptors as initially shown in neonatal neurons of the facial nucleus in the brainstem (Raggenbass et al., 1991). Recent data also show a direct increase in neuronal excitability by AVP via suppression of K^+^-channels in the hippocampal formation including subiculum (Lei et al., 2021), CA1 (Hu et al., 2022) and dentate gyrus (Lei et al., 2022) as well as in the medial aspect of the central amygdala (Boyle et al., 2021). V_1a_ receptors are coupled to Gα_q/11_ proteins activating phospholipase C_β_ (PLC_β_), which hydrolyses phosphatidylinositol 4-5-biphosphate (PIP_2_) to inositol triphosphate (IP_3_) and diacylglycerol (DAG). In turn, IP3 leads to increased Ca^2+^ release from intracellular stores and DAG activated protein kinase C (PKC). In CA1 of young rats, direct excitation of pyramidal neurons by AVP involves a persistent inward current and is mediated by V_1a_ receptors leading to suppression of G protein-gated inwardly rectifying K^+^ (GIRK) channels via PLC_β_-induced degradation of PIP_2_ (Hu et al., 2022). Increase in intracellular Ca^2+^-release and PKC activation were not involved in the effect. GIRK function is not detectable in neonatal mouse 5-HT neurons, and it develops between P4-P21 to adult-like levels (Rood et al., 2014). Hence, a potential mechanism for the direct excitation of 5-HT neurons by AVP at P10-12 in our experiments could be V_1a_ mediated suppression of GIRK channels which may become even more pronounced with further maturation of GIRK function. While about 20% of 5-HT neurons in adult mouse DRN express V_1a_ receptors, only a small fraction (10%) of V_1a_ receptor expressing neurons contains 5-HT (Patel et al., 2022), suggesting that in the adult rodent DRN AVP mostly signals to non-serotonergic neurons. However, this may be distinct in the developing DRN. It is well established that there is a transient increased expression of V_1a_ receptors in certain brain regions depending on the species during postnatal development in rodents which peaks around the end of the first postnatal week and then declines until adulthood (Hammock, 2015). In rats, AVP receptor expression in DRN is much stronger at P10 than in adulthood (Tribollet et al., 1991). Hence, it is likely that during neonatal development a higher proportion of 5-HT neurons expresses V_1a_ receptors than in adulthood. This may explain why direct excitation of 5-HT neurons by AVP is frequent during postnatal development but rare during adulthood.

### AVP does not affect inhibitory synaptic transmission in the neonatal DRN

While bath application of AVP caused a prominent increase in the frequency of sEPSCs with or without blocking GABA_A_ receptors, neither sIPSCs frequency nor amplitude were altered by AVP. This is in line with data from adult mouse DRN where AVP did not increase sIPSC frequency either (Rood and Beck, 2014). However, recent transcriptome profiling and in situ hybridization identified at least three distinct populations of V_1a_ receptor-expressing neurons in DRN, including GABAergic neurons (Patel et al., 2022), suggesting that they also do respond to AVP under certain conditions and modulate the predominant glutamatergic excitatory response. Another reason why sIPSCs in 5-HT neurons are not altered by AVP in neonatal DRN may be the protracted development of GABAergic transmission in DRN. Similar to our recordings from rat neonatal 5-HT neurons, the frequency of baseline sIPSCs in neonatal mouse DRN is very low during early postnatal development and dramatically increases until adulthood (Rood et al., 2014; Morton et al., 2015). A steep increase in GABAergic synapses onto 5-HT neurons has been shown between P4-21 (Arganaraz et al., 2022), that may underlie the increase in GABAergic synaptic inputs. Other contributing factors could be an increase in spontaneous firing activity of GABAergic neurons in DRN or changes in GABA release probability (Morton et al., 2015). Hence, whether a direct response of V_1a_ receptor expressing neonatal GABAergic neurons to AVP will be signalled to 5-HT neurons may strongly depend on the functional maturation of GABAergic synaptic transmission in DRN.

### Neonatal 5-HT neurons are characterized by slow oscillating firing as well as tonic regular activity *in vivo*

5-HT neurons in adult anaesthetised rodents are characterized by their tonic clock-like firing pattern at low rate (Allers and Sharp, 2003; Andrade et al., 2015) as well as broad AP waveform with long afterhyperpolarization. This clock-like firing activity and broad AP waveform was also common in identified 5-HT neurons in the neonatal DRN. However, the largest proportion of identified 5-HT neurons we encountered *in vivo* in neonatal DRN had low frequency oscillations in firing activity. To our knowledge, this firing pattern of DRN neurons has only been described in two early studies, one in brain slices and one in chloral hydrate anaesthetised adult rats (Mosko and Jacobs, 1974, 1976) as well as in one more recent *in vitro* electrophysiological characterization of genetically identified 5-HT neurons in mouse DRN (Mlinar et al., 2016). In this study, carried out in 4–28-week-old mice, approx. 1% of recorded 5-HT neurons exhibited a LFO in firing pattern. The broad range of the period of oscillations from 10 to up to 60 s as well as the described intermittent occurrence of spike trains with up to 40s silent period in between is very similar to the *in vivo* LFO firing of neonatal 5-HT neurons in our study. Furthermore, they also describe transitions of the same cell between tonic regular and slow oscillating firing as we have also observed. Hence, we can confirm that slow oscillation firing is indeed an alternative firing mode rather than an attribute of a distinct population of 5-HT neurons. The fact that hardly any studies have encountered this slow oscillating firing pattern during adulthood while in our recordings this was predominant suggests that this is a signature of developing 5-HT neurons. It is intriguing, that mice used by Mlinar et al. 2016 also included 4 week old animals which is just the age when the function of 5-HT neurons starts to become adult-like (Rood et al., 2014).Hence the presence of slow oscillating firing in their sample may be related to immature 5-HT neurons. What determines this firing mode, whether this is driven by a specific type of input predominant during development, an immature balance of different afferent synaptic inputs that are still changing in strength or even an intrinsic oscillation mode due to immature expression of ion channels or receptors on 5-HT neurons will be interesting to investigate in future studies.

### AVP also excites putative DRN 5-HT neurons *in vivo*

We tested here for the first time the effect of AVP on the firing activity of DRN neurons *in vivo* and found that the majority of neurons were responding to AVP with either increase or decrease in firing activity. Since our multi-unit recordings included all types of active neurons in DRN which encompass a heterogeneous population (Cardozo Pinto et al., 2019), not only 5-HT neurons, this may represent the response of different cell types in DRN expressing the V_1a_ receptor. However, while vehicle injections did not affect the firing activity of the majority of neurons, about one fourth (26%) were inhibited by vehicle injections which may be a technical artefact of the injection in close proximity to the recording site. Accordingly, the inhibitory response of 35% of DRN neurons to AVP may also be an artefact of the injection and has to be interpreted with caution. Future experiments need to confirm the inhibitory effect of AVP *in vivo* by driving internal release. The largest proportion of neurons though were excited by AVP which is in line with our *in vitro* findings and reports that predominantly glutamate containing neurons in DRN express V_1a_ receptors (Patel et al., 2022) that may relay this response to 5-HT neurons. The majority of activated neurons showed similar tonic regular firing or a pattern of low frequency oscillations as we had found is characteristic in a smaller sample of identified neonatal 5-HT neurons. Since activated neurons also had a long afterhyperpolarization as typical for neonatal 5-HT neurons, this strongly suggests that 5-HT neurons are also activated by AVP *in vivo*.

### The vasopressinergic innervation of neonatal DRN is still sparse and originates in medial amygdala and BNST

We demonstrated here that in the DRN of the neonatal rat the vasopressinergic innervation is already present but still sparse in comparison to data in adult animals (Rood and De Vries, 2011). However, typical features, such as its strong sex dependence and dorsal to ventral gradient described in adulthood (Rood and De Vries, 2011; Rood et al., 2013) are already present during postnatal development. The presence of vasopressinergic innervation in DRN at a developmental time point corresponding to human birth provides the anatomical substrate which may mediate increased activation and release of 5-HT during complicated birth through strong activation of the central AVP system. However, if this neonatal sparse innervation is fully functional and contains active release sites has to be tested in the future by driving internal release by e.g. optogenetic or chemogenetic stimulation of immature AVP neurons in BNST or medial amygdala.

Since the vasopressinergic innervation of DRN is dependent on gonadal steroid hormones, the sparse innervation during neonatal development may reflect the low levels of circulating sex hormones at this age, which, nonetheless, are sufficient to maintain some degree of AVP-immunoreactivity in males. This suggest that the vasopressinergic innervation of the DRN undergoes developmental changes which is likely driven by the rise in circulating gonadal steroid hormones during puberty. Developmental changes, i.e. an increase in strength in the vasopressinergic innervation between juvenile development and adulthood as well as sex differences have also been shown in other brain regions receiving vasopressinergic innervation from the BNST and medial amygdala including the lateral septum (DiBenedictis et al., 2017). While the function of AVP in adult DRN is still hardly understood, it has been shown to play a role in some social behaviors such as response to a female social stimulus in both males and females (Patel et al., 2022) or urine marking in males towards unfamiliar males but not females (Rigney et al., 2020). It is likely that the age-dependent increase in vasopressinergic innervation of DRN, resulting availability of AVP and control over DRN network function and output may serve to regulate age-specific social behaviors. Future studies are necessary to better understand the role of AVP for behaviors governed by DRN and the particular role of the developmental increase in innervation/availability of AVP therein.

To our knowledge we conducted the first retrograde tracing study to investigate the origin of the vasopressinergic innervation of DRN. Our findings that this innervation originates in cell groups in BNST and medial amygdala is in line with previous studies using gonadectomy that suggested that DRN receives vasopressin fibers originating from cell groups in BNST and/or medial amygdala in adult mice (Rood et al., 2013). This was then directly confirmed by recent anterograde viral tracing experiments in adult mice showing that both BNST and medial amygdala AVP neurons project to DRN (Rigney et al., 2023). Here we confirm this fiber origin in another species, the rat, and show that vasopressinergic cell groups in both regions, BNST and medial amygdala, project to DRN already during neonatal development. We found that in neonatal males, both BNST and medial amygdala contained similar numbers of AVP neurons and that a similar ratio of these cells were projecting to DRN, suggesting that they contribute equally to the early vasopressinergic innervation of DRN. This may change during following development as the number of anterogradely labelled synaptophysin puncta in DRN originating from AVP neurons in BNST was larger than from AVP neurons originating in medial amygdala (Rigney et al., 2023) in adult mice suggesting that the innervation from BNST AVP neurons may become more predominant than from medial amygdala AVP neurons in adulthood. We also showed that the vasopressinergic innervation from BNST and medial amygdala is exclusive and that AVP neurons in other brain regions such as PVN do not contribute to the innervation, at least during development.

To date it is not known whether only hypothalamic or also extra-hypothalamic AVP neurons are activated during birth. A recent study in both mice and rats showed a strong increase in cFOS positive (+) neurons after vaginal birth as an indirect marker for neuronal activation in SCN, SON and PVN of the hypothalamus in either species which were mainly vasopressinergic (Hoffiz et al., 2021). The same study also reported an increase in cFOS^+^ neurons in medial amygdala but did not investigate their neurochemical phenotype. Hence, it is possible that AVP neurons in medial amygdala and BNST are also directly activated during labour which needs to be investigated in future studies. Interestingly, the authors also showed that premature birth led to an even more widespread activation of AVP neurons in PVN that they explain with increased activation by greater hypoxia during premature delivery (Kamlin et al., 2006; Dawson et al., 2010) than during full-term delivery. Since birth stress is strongly associated with prolonged hypoxia and an exaggerated surge of AVP in the periphery (Schlapbach et al., 2011; Evers and Wellmann, 2016; Summanen et al., 2018; Fill Malfertheiner et al., 2021), this may involve increased activation of AVP neurons both in intensity and number in PVN. AVP neurons in BNST and medial amygdala are widely interconnected with each other but also receive inputs from AVP neurons in PVN (Rigney et al., 2023). It is tempting to speculate that greater activation of AVP neurons in PVN by prolonged hypoxia during birth stress may cause additional activation of BNST and medial amygdala neurons through direct afferents from AVP neurons in PVN. The either direct and/or indirect activation of BNST and medial amygdala AVP neurons by prolonged hypoxia during birth stress may results in increased AVP release in target regions such as the DRN causing strong activation of neonatal 5-HT neurons.

### What is the role of AVP in neonatal brain networks?

It is well established that vaginal birth leads to a surge of AVP in the blood that promotes survival and adaptation of the fetus to extra-uterine life by facilitating air breathing and the adjustment of blood pressure, maintenance of body temperature as well as glucose and water homeostasis (Hillman et al., 2012). Furthermore, it is likely that central levels of AVP in the brain are also increased during and after labour, since neonates also showed elevated AVP levels in CSF which correlated well with serum levels (Bartrons et al., 1993; Carson et al., 2014). To investigate the role of AVP in the perinatal brain, Spoljaric et al. 2017 tested the effect of AVP in the neonatal hippocampus using *in vitro* electrophysiological methods. They found that AVP powerfully increased inhibitory GABAergic synaptic transmission onto pyramidal neurons by enhacing the firing activity of GABAergic interneurons. This suggested that AVP is protective in the perinatal brain which would be in line with its survival promoting effects in the periphery. Our results from neonatal rat DRN however show a prominent excitatory effect of AVP during a period equivalent to human birth, suggesting that the effect of AVP on perinatal subcortical networks may be distinct from its effect in hippocampal and cortical networks. Both hypothalamic and extra-hypothalamic AVP neurons have widespread projection targets in the brain, the former including both cortical and subcortical brain regions (Rood et al., 2013) while the latter has predominantly subcortical targets with exception of the dorsolateral entorhinal cortex and ventral CA1 in hippocampus (Rigney et al., 2023). Subcortical brain regions particularly in the hindbrain, such as the solitary nucleus or lateral parabrachial nucleus, innervated by hypothalamic AVP neurons in PVN are involved in the regulation of the internal physiological state such as blood pressure, water balance and stress response (Andresen and Kunze, 1994; Menani et al., 1996; Rood et al., 2013). Since such regulation is critical during the transition from uterine to extra-uterine life, it is unlikely that AVP would be predominantly inhibitory in those networks at the time of birth as it has been shown for hippocampal networks. Rather, the effect may differ between brain regions depending on their function during birth, the expression profile of V_1a_ receptors within the local network during development or the maturation of GABAergic synaptic transmission. It is likely that there is a fine balance between inhibitory and excitatory effects of AVP in the brain at birth that supports the transition to extra-uterine life and promotes survival of the neonate. Hyperactivation of the perinatal AVP system by prolonged hypoxia during complicated birth or mild birth asphyxia may cause this balance to break down with lasting effects on the development of immature networks and an outcome that renders more susceptible to disease later on.

### Functional considerations of strong activation of the neonatal 5-HT system by AVP at the time of birth

It is well established that 5-HT plays a strong role during brain development where it not only acts as a neurotransmitter but also as a trophic factor (Gaspar et al., 2003). Furthermore, pharmacologically elevated 5-HT levels during early postnatal development spanning approx. the first two postnatal weeks of rodent development, also termed the 5-HT sensitive period, promote increased anxiety and depressive-like behaviors in rodents (Ansorge et al., 2004; Rebello et al., 2014). Furthermore, an association of prenatal exposure to selective serotonin re-uptake inhibitors (SSRIs) that elevate 5-HT levels with increased risk for depression in adolescence has also been shown in humans (Malm et al., 2016). DRN 5-HT neurons have a protracted functional development that is characterised by hyperexcitability, absence of 5-HT_1a_ autoreceptor control and lack of GABAergic synaptic input during early postnatal development that only become adult-like after 3-4 weeks postnatally in mice (Rood et al., 2014). Furthermore, significant refinement of glutamatergic synapses in strength (Kisner and Polter, 2023) but also in number particularly from cortical inputs onto developing 5-HT neurons in DRN (Arganaraz et al., 2022) occurs during a similar postnatal period. Together this suggests that the first 3-4 postnatal weeks in rodents comprise a period of heightened plasticity for 5-HT neurons which renders them extremely sensitive to environmental perturbations during this time. As rodents are born immature, the equivalent time point of human birth lies during rodent postnatal development at around P10-12 in rats (Workman et al., 2013). Strong activation of immature 5-HT neurons by increased AVP release triggered by prolonged periods of hypoxia during complicated labour or mild birth asphyxia may affect their own ongoing functional maturation during this sensitive period of heightened plasticity or may even lead to excitotoxicity. Furthermore, persistent changes in their activity may also result in an imbalance in early 5-HT levels in target regions that could affect the maturation of those networks as demonstrated e.g. for mPFC (Rebello et al., 2014) and contribute to increased risk to psychiatric disorders later on. Further studies are needed to directly test the effect of birth asphyxia on the development of the 5-HT system and the role of AVP therein in rodent models.

### Conclusions

This work clearly demonstrated that nanomolar concentrations of AVP strongly excite neonatal 5-HT neurons in DRN in a V_1a_ receptor dependent manner and lead to an increase in action potential firing both *in vitro* and *in vivo.* This is mediated by a combination of an increase in glutamatergic synaptic input and most likely also a direct mechanism onto those neurons while GABAergic synaptic inputs were not affected. We also show that some degree of vasopressinergic innervation is already present in neonatal DRN in a sex dependent manner in males that may form the anatomical substrate for the excitatory effect of AVP. Furthermore, we confirm that the vasopressinergic innervation of DRN originates from AVP neurons in BNST and medial amygdala, and show that it is exclusive with approximately equal contribution from either region. While it has previously been shown that 5-HT neurons undergo a protracted functional development *in vitro* (Rood et al., 2014), we provide the first characterisation of firing activity of 5-HT neurons during neonatal development *in vivo*. In addition to the typical tonic clock-like firing pattern that are typical for 5-HT neurons in adulthood, we found that a large proportion of neonatal 5-HT neurons shows low frequency oscillations in firing, suggesting that also their *in vivo* firing activity undergoes a protracted postnatal development. While it has been shown that AVP inhibits neonatal hippocampal networks and hence has been termed protective during birth, our data show that the effect of AVP in subcortical networks such as the DRN is distinct from that and in contrast highly excitable. Strong activation of the neonatal 5-HT system by AVP during birth stress may affect ongoing brain development. Future studies further investigating the role of AVP in both cortical and subcortical networks at the time of birth will be valuable in expanding our knowledge on its early function.

## METHODS

### Animals

Experiments were performed using Wistar rat pups of both sexes at ages defined in each experiment ranging from postnatal day (P) 10 to 16. All experiments with animals were carried out in accordance with University of Helsinki Animal Welfare Guidelines and approved by the Animal Experiment Board in Finland (license ESAVI/13422/2018 and ESAVI/27422/2022).

### *In vitro* electrophysiology

*Acute slice preparation.* 300 µm coronal acute slices were prepared from brains of 10 – 12-day-old male and female Wistar rat pups. Solutions used during slice preparation and recording were adopted from Rood et al., 2014. During the dissection and slicing the brain was kept in an ice-cold solution containing (in mM): 248 Sucrose, 2.5 KCl, 1.25 NaH_2_PO_4_, 2 MgSO_4_, 2.5 CaCl_2_, 10 dextrose, 26 NaHCO_3_ (all from Sigma-Aldrich) and saturated with 95% O_2_ and 5% CO_2_. Immediately after cutting the slices were transferred to ACSF containing (in mM): 124 NaCl, 2.5 KCl, 1.25 NaH_2_PO_4_, 2 MgSO_4_, 26 NaHCO_3_, 10 D-glucose, 2.5 CaCl_2_, 2.5-5 L-tryptophan; 5 % CO_2_ / 95% O_2_ and incubated 60 min at +35°C and up to 4 h at room temperature before use. The slice was placed in a submerged recording chamber and continuously perfused with ACSF without L-tryptophan at +30°C.

*Electrophysiological recordings.* Whole-cell voltage clamp recordings were made from DRN under visual guidance using patch electrodes with resistance of 4–6 MΩ. Uncompensated series resistance (Rs) was monitored by measuring the peak amplitude of the current response to a 5-mV step. Only experiments where Rs < 30 MΩ, and with < 20% change in Rs during the experiment, were included in the analysis.

*Spontaneous excitatory (sEPSC) and inhibitory (sIPSC) postsynaptic currents* were recorded simultaneously using patch pipettes filled with a solution containing (in mM): 135 K-gluconate, 10 HEPES, 2 KCl, 2 Ca(OH)_2_, 5 EGTA, 4 Mg-ATP, 0.5 Na-GTP; 285 mOsm and pH was adjusted to 7.2 with NaOH (all from Sigma-Aldrich). sEPSC/sIPSC were recorded at a holding potential of –47 mV.

*Pharmacologically isolated sEPSC* were recorded using a patch pipette filled with a solution containing (in mM): 120 CsMeSO_4_, 5 NaCl, 10 TEA-Cl, 10 HEPES, 5 QX-314 chloride, 0.5 EGTA, 4 Mg-ATP, 0.3 Na-GTP; 280 mOsm and pH was adjusted to 7.2 with CsOH (all from Sigma-Aldrich). sEPSCs were recorded at a holding potential of –70 mV in the presence of 100 uM Picrotoxin to block GABAergic transmission via GABA_A_ receptors.

*Spontaneous spiking activity* was recorded in cell attached mode using 6-8 MΩ recording pipette filled with 150 mM NaCl (see Perkins, 2006).

*Pharmacology.* Table 1 depicts all pharmacological tools used in *in vitro* electrophysiology experiments in this study.

**Table 1.** Pharmacological tools used in this study.

| Drug | Function | Working concentration | Manufacturer |
| --- | --- | --- | --- |
| [Arg<sup>8</sup>]-Vasopressin (AVP) | Vasopressin receptor agonist | 10 nM | Tocris |
| L-tryptophan | Precursor of serotonin | 2.5 – 5 μM | Sigma-Aldrich |
| SR49059 | V<sub>1a</sub> antagonist | 20 nM | Tocris |
| SSR149415 | V<sub>1b</sub> antagonist | 20 nM | Tocris |
| TGOT | Oxytocin receptor agonist | 10 nM | Bachem |
| Picrotoxin | GABA<sub>A</sub> antagonist | 100 μM | Abcam |
| D-AP5 | NMDA antagonist | 50 μM | HelloBio, UK |
| CNQX | AMPA/KAR antagonist | 10 μM | HelloBio, UK |
| MCPG | group I/ II mGluR antagonist | 250 μM | HelloBio, UK |
| CGP55845 | GABA<sub>B</sub> antagonist | 10 μM | Tocris |
| WB4101 | Alpha1 adrenergic receptor antagonist | 1 μM | Sigma-Aldrich |

**Table 2.** Mean number of double positive (double+), Neurophysin-2 positive (Neurophysin-2+) and FG positive (FG+) neurons in MeAmy, BNST, PVN and SON of N=5 rat pups.

|  | double+ neurons | Neurophysin-2+ neurons | FG+ neurons |
| --- | --- | --- | --- |
| MeAmy | $58.00 \pm 11.81$ | $224.8 \pm 52.67$ | $469.20 \pm 132.03$ |
| BNST | $38.60 \pm 4.93$ | $207.80 \pm 41.03$ | $964.6 \pm 220.61$ |
| PVN | $1.4 \pm 1.17$ | $633.00 \pm 136.56$ | $426.00 \pm 118.96$ |
| SON | $1.2 \pm 0.97$ | $973.6 \pm 372.63$ | $43.2 \pm 33.50$ |

*Biocytin filling of neurons.* Biocytin (0.1%) (Sigma-Aldrich) was added to intracellular solutions and pipette solution in cell attached mode to later visualize and analyse cell morphology and to confirm the location of the recorded neuron. Neurons recorded in cell attached mode were filled after the recording by clamping the neuron at holding potential –70 mV and going whole-cell. After filling the recorded neuron with biocytin, the slice was fixed with 4% paraformaldehyde (PFA) in PBS.

*Immunohistochemistry.* Anti-TPH and streptavidin double labelling was used to identify TPH positive serotonergic neurons from all recorded biocytin filled neurons. 300 µm thick fixed sections collected after electrophysiology recordings were washed using PBS and PBS + 0.3% Triton X-100 (PBST) for 5 x 7 min and incubated 2h at room temperature (RT) with blocking solution containing 10% normal donkey serum (NDS, Merck Millipore), 3% Bovine serum albumin (BSA, Sigma-Aldrich) in PBST. Sections were then incubated 48 – 72 h with sheep anti TPH (1:1000, ab1541, Merck Millipore) primary antibody in carrier solution containing 1% NGS and 1% BSA in PBST at +4°C. Sections were washed 5 x 7 min with PBS and PBST, and incubated over-night at +4°C with donkey anti-sheep secondary antibody conjugated with Alexa Fluor 568 (1:1000, A-11011, Invitrogen, ThermoFisher Scientific) and Streptavidin conjugated with Alexa Fluor 488 (1:2000) in carrier solution. Sections were rinsed 5 x 7 min with PBS, and mounted using Fluoromount-G (00-4958-02, Invitrogen, ThermoFisher Scientific).

*Imaging.* Brain sections were imaged using a Zeiss fluorescent microscope with Apotome (Axio Imager 2, Carl Zeiss, Oberkochen, Germany) and processed and exported with the Zeiss Zen Lite Software (Carl Zeiss, Oberkochen, Germany).

*Data analysis.* Data were collected using WinLTP software (WinLTP Ltd, Bristol, UK) and analysed offline using the MiniAnalysis 6.0.3 program (Synaptosoft Inc., Fort Lee, NJ, USA). The threshold for detection of both inward (sEPSC, action potentials) and outward (sIPSCs) events was three times the baseline noise level, and all detected events were verified visually. For comparison and quantification of drug effects (see bar diagrams), events were extracted into 10 s bins to determine an appropriate analysis period around the peak effect. This period was then chosen for analysis using the raw data. Accordingly, baseline activity was quantified during the 3 min preceding drug application. Here the start of the drug application refers to the change in solution of the inlet tube, i.e. includes the time the drug solution needs to travel to the recording chamber. The effect of AVP or TGOT on sEPSC frequency and amplitude was calculated for a period of 60 s between 190-250 s after the beginning of drug application. The same interval in relation to AVP application was also quantified during co-application with receptor blockers. The effect of AVP alone or in the presence of glutamatergic or GABAergic receptor blockers on action potential firing was analysed for a period of 30 s from 270-300 s after beginning of drug application. In the combined sEPSC/sIPSCrecordings, sEPSC frequency and amplitude was analysed for a period of 90 s from 420-510 s after the start of drug application. sIPSC frequency and amplitude was analysed for 110 s from 180-290 s after the start of the drug application.

### Neurophyisin2 fiber fractional area analysis in DRN

*Tissue preparation and immunohistochemistry.* In total 14 neonatal Wistar rat pups (seven male, seven female) were deeply anaesthetised with pentobarbital and then transcardially perfused with 4% PFA dissolved in 0.1 M borate buffer. Brains were removed and post-fixed in 4% PFA for 1-2 days at 4°C. Free-floating 70 µm coronal sections of the DRN were cut using a vibratome (Leica VT1200). Sections were washed 3 x 7 min in PBS and incubated 1h with blocking solution containing 10% normal goat serum (NGS, MP Biomedicals, USA), 3% BSA and 0,3% Triton-X in PBS. Sections were then incubated for 24 hrs with rabbit anti-5-HT (AB572263, 1:1000, Immunostar) and mouse anti-Neurophysin2 (PS-41, 1:100, generously provided by Dr. Harold Gainer, National Institute of Neurological Disorders and Stroke, NIH, USA) primary antibodies in carrier solution containing 1% NGS, 1% BSA and 0,3% Triton-X in PBS. Sections were washed 3 x 7 min with PBS and incubated overnight at +4°C with goat anti-rabbit Alexa Fluor 594 and goat anti-mouse Alexa Fluor 488 (each 1:1000, Invitrogen, ThermoFisher Scientific) in carrier solution. Sections were washed 3 x 7 min with PBS and mounted with Fluoromount-G.

*Light microscopy imaging.* Light microscopy images from brain sections containing DRN were obtained with the Axio Imager.M2 microscope equipped with an Apotome 2 structured illumination microscopy slider, using EC Plan-Neofluar 5x (NA = 0.16), PlanApo oil-immersion 40x (NA = 1.30) objectives (all from Zeiss), fitted with a Hamamatsu ORCA Flash 4.0 V2 camera. The ZEN 2 software (Zeiss) was used for image acquisition. Images with 5x objective were obtained to determine the area and outline of DRN according to the distribution of 5-HT-immunoreactive (ir) neurons. Based on evaluation of 5x images, one section at similar anatomical levels of rostral, central and caudal DRN was chosen from each pup for further z-stack imaging at 40x. From rostral DRN 5-6, from central DRN 8 and from caudal DRN 3-4 40x stack images were obtained covering the whole DRN area in each section.

### Quantitive fiber fractional area analysis

AVP fiber fractional area (FFA) analysis was performed similarly as described by DiBenedictis et al., 2017 using Fiji/ImageJ software (version1.51 u). First, the maximum intensity projection was performed on each 40x z-stack image and the resulting image was adjusted for brightness and contrast. Then manual thresholding of the gray-scale image was performed by adjusting the threshold so that the background would be clear white and the fiber signal black. Artefacts were removed from the image using a freehand selection tool and “fill” command. Finally, the FFA percentage (% of black pixels) was measured.

For comparison of AVP FFAP between rostral, central and caudal DRN, values from all images from each pup at the respective DRN level were pooled. For comparison of AVP FFAP between dorsal and ventral DRN, values from 4 images covering dorsal DRN at rostral levels, 6 at central and 2 at caudal levels were pooled. Accordingly, values from 2 images covering ventral DRN at rostral levels, 2 at central and 2 at caudal levels were pooled. For comparison of AVP FFAP in males vs females, all values from each pup were averaged.

### Anatomical tract tracing with the retrograde tracer Fluorogold

*Surgical preparation.* The retrograde tracer Fluorogold (FG, Fluorochrome, USA) was delivered into DRN of male P12 Wistar rat pups by iontophoresis. For this, anaesthesia was induced by isoflurane (4%) and pups were injected with Carprofen i.p. (Rimadyl, 1% solution, 5µg/g) for analgesia before starting the surgery. Pups were transferred into a stereotactic frame and the head fixed in place. Body temperature was kept constant at 37° C by a biological temperature controller (Supertech Instruments, UK). Isoflurane concentration was reduced to 1-2 % for the anaesthesia maintenance. Lidocaine for local anaesthesia was injected subcutaneously below the skin above the area of interest. A small incision was made only exposing the skull above λ. A burr hole was drilled 0.4-0.5 mm behind λ exposing the central sinus. A glass pipette (40 µm tip) filled with 2% FG was lowered into DRN targeting the middle of the sinus at depth 5.2 mm from the brain surface. FG was expelled for 5 min using an intermittent protocol (7s on, 5s off, +/ – 7 µA). The pipette was then retracted 200 µm and FG expelled for 10 min using the same protocol as before. This ensured a sufficient size of the injection site spanning the entire DRN region. After the iontophoresis, the pipette was left in place a further 5 min and was then retracted. The wound was closed with tissue glue (Vetbond, 3M). Animals were transferred to a clean cage on a heating pad for recovery from anaesthesia and were then returned to the dam. Pups wellbeing and body weight was controlled daily and additional carprofen doses as above were also administered daily. Four days after the FG delivery, pups were deeply anaesthetised at P16 with pentobarbital and the brains fixed by transcardial perfusion with 4% PFA in PBS.

*Immunohistochemistry.* After 1-3 days postfixation in 4% PFA at 4°C, free-floating 70 µm coronal sections of the DRN were cut using a vibratome (Leica VT1200). Sections were washed 3 x 10 min in PBS and incubated 1h with blocking solution containing 10% normal goat serum (NGS, MP Biomedicals, USA), 3% BSA in PBST. Sections were then incubated 24 – 48 h with rabbit anti-fluorescent gold (1:1000, AB153-I, Millipore) and mouse anti-Neurophysin2 (PS-41, 1:100, generously provided by Dr. Harold Gainer, National Institute of Neurological Disorders and Stroke, NIH, USA) primary antibodies in carrier solution containing 1% NGS, 1% BSA in PBST. Sections were washed 3 x 10 min with PBS and incubated over-night at +4°C with goat anti-rabbit Alexa Fluor 564 (1:1000, Invitrogen, ThermoFisher Scientific) and goat anti-mouse Alexa Fluor 488 (1:1000, Invitrogen, ThermoFisher Scientific). Sections were washed 3 x 10 min with PBS and mounted with Fluoromount-G.

*Image analysis.* Brain sections containing the Medial Amygdala (MeAmygd), Bed Nucleus of Stria Terminalis (BNST), Paraventricular nucleus of the Hypothalamus (PVN) and Supraoptic nucleus of the Hypothalamus (SON) were imaged as described above. To quantify AVPergic neurons projecting to DRN, Fluorogold and Neurophysin2 double-positive neurons were counted and normalized to total number of Neurophysin2 positive neurons in each region. All sections that contained Neurophysin2 positive neurons in these brain regions were subjected to data quantification. ImageJ software was used to count Fluorogold and Neurophysin2 positive neurons, labelled neurons were verified visually.

### *In vivo* electrophysiology

*Surgical preparation.* Extracellular recordings were performed in dorsal raphe nucleus (DRN) of postnatal day (P)10-12 male and female rats. Anaesthesia was induced with isoflurane (4%) followed by i.p. administration of urethane (1g/kg; Sigma-Aldrich). Isoflurane administration (1-2%) continued during surgery to ensure deep anaesthesia. A small burr hole was drilled above the region of interest (DRN: 6.5-7.0 mm posterior to bregma on the midline) and the bone was carefully removed. The underlying dura mater was thinned using forceps exposing the central sinus. A small drop of mineral oil was applied to the burr hole to keep the underlying tissue moist. One additional burr hole each was drilled above the left frontal cortex and contralateral cerebellum serving as reference for recording of electrocorticograms (high frequency-pass filtered at 0.1 Hz with x2000 amplification) via steel skull screws to monitor the depth of anesthesia. The head of the pup was fixed into the stereotaxic apparatus (Stoelting, Wood Dale, IL) using two plastic bars fixed with dental cement on the nasal and occipital bones, respectively. The body temperature was maintained at 37 °C with a thermoregulated heating blanket (Supertech Instruments, UK). A urethane top-up of 1/3 of the original dose was administered if needed.

*Juxtacellular single unit recording and labelling.* A glass recording microelectrode (12-25 MΩ impedance *in vivo*) filled with 0.5 M NaCl and neurobiotin (1.5% w/v; Vector Laboratories, UK) was lowered into the DRN using a motorized manipulator (IVM-1000, Scientifica, UK) to a depth of 4.5-5.2 mm from the surface. Biopotentials were amplified (x10) through the active bridge circuitry of a Neurodata amplifier (Cygnus Technologies, USA), AC-coupled and further amplified (x100; NL106 AC-DC Amp; NeuroLog System, Digitimer Ltd, UK) before being filtered between 0.3 and 5 kHz (NL125/NL126 filters; NeuroLoig System, Digitimer Ltd, UK). Signals were digitized online using a Micro1401 Analog-Digital converter (Cambridge Electronic Design (CED), UK) and a PC running Spike2v9 acquisition and analysis software (CED, UK). Single unit activity and electrocorticograms were sampled at 20 kHz. After recording the baseline activity of a spontaneously active DRN neuron for 5 min, the neuron was entrained for 30 s to 5 min using 1-10 nA 200 ms on/off current pulses passed through the recording electrode. In 19 pups, a single neuron was labelled, in seven animals two neurons were labelled and in two animals three neurons were labelled. In the case of multi-labelling, neurons could be unambiguously identified due to the large difference in recording location. After labelling, pups were maintained under anaesthesia for a further 0.5-2 hours followed by perfusion/fixation (see below).

*Multi-unit activity recording and in vivo drug application in DRN.* A multielectrode array (Cambridge Neurotech, UK) was inserted perpendicularly to the skull surface into DRN until a depth of 4.8-5.2 mm. The electrodes were labeled with DiI (1,1’-dioctadecyl-3,3,3’,3’-tetramethyl indocarbocyanine, Invitrogen) to enable reconstruction of the electrode tracks post mortem (Fig. 5A). A silver wire inserted into the cerebellum served as ground and reference electrode. After 15-20 min recovery period, simultaneous recordings of multi-unit activity (MUA) and local field potential (LFP) were performed from the DRN using single shank 64-channel silicon probes (H9, ∼50 kΩ). The recording sites were separated by 45 µm. The position of recording sites within DRN was confirmed by post-mortem histological evaluation.

For local injection of either vehicle or drug into DRN, a glass pipette with a tip diameter of 30-50 µm filled with either vehicle (0.9% NaCl) or 100 nM AVP was lowered 0-100 µm behind the recording probe at an angle of 10° into the DRN (mediolateral 0.8 mm from sinus midline, dorsoventral 4.7-4.9 mm from the sinus surface).

Both LFP and MUA were recorded at a sampling rate of 32 or 30 kHz using a multi-channel extracellular amplifier (Digital Lynx 4SX, Neuralynx, USA or Open Ephys acquisition board, Open Ephys, USA) and the corresponding acquisition software. During recording the signal was band-pass filtered between 0.1 Hz and 8 kHz (Neuralynx) or 0.1 Hz and 7.9 kHz (Open Ephys). After 15 min of baseline recording, 200 nl of either vehicle or AVP were injected at a speed of 70 nl/ min using a microinjection syringe pump (UMP3, World Precision Instruments, USA) followed by a further 15 min of baseline recording afterwards.

*Perfusion/Fixation.* After the recording, pups received another dose of more concentrated urethane (2g/ kg) for deep surgical anaesthesia and were perfused with 0.9% NaCl followed by 4% paraformaldehyde. Brains were postfixed in 4% PFA for a further 3-5 days.

*Immunohistochemistry.* Free-floating 70 µm coronal sections containing DRN were cut on a vibratome. Sections were washed 3 x 10 min in PBS and incubated for 1h in blocking solution containing 10% NGS and 3% BSA in PBST at RT. Sections were incubated for 24-72 hrs at RT with rabbit anti-5-HT primary antibody (1:1000, cat no 20080, Immunostar, USA) in carrier solution containing 1% NGS, 1% BSA in PBST. Sections were rinsed 3x 10 min with PBS and incubated either for 2 hrs at RT or 24-72 hrs at 4° C in goat anti-rabbit Alexa Fluor 488 (1:1000, Invitrogen, ThermoFisher Scientific) secondary antibody in carrier solution containing 1% NGS, 1% BSA in PBST. To reveal the Neurobiotin filled neurons, streptavidin 594 (1:2000, Invitrogen, ThermoFisher Scientific) was added to the secondary antibody solution. Sections were washed 3 x 10 min with PBS and mounted with Fluoromount-G. Sections were imaged as described above.

*Data analysis*. To isolate single units from multi-unit activity (MUA) recordings, data were imported into Plexon Offline sorter version 4.4.2 (Plexon Instruments, USA). Raw signals from each channel were high-pass filtered above 400 Hz (Butterworth, 2^nd^ order high-pass filter) and spikes were detected by using amplitude threshold that was set manually after visual inspection of the signal-to-noise ratio (SNR) in each channel. The detected spikes were automatically sorted into individual units using the valley-seeking method, a clustering algorithm that identifies the cluster centres of the principal components calculated from the derivatives of the action potential waveforms. Units with a SNR below five were excluded. Individual spiking units were identified if their corresponding spike clusters were clearly separated in feature space, the waveform shape was consistent with a clear resemblance to derivatives of action potentials and the interspike interval (ISI) histogram showed a refractory period of >1 ms (Plexon Offline sorter). If a unit occurred simultaneously in more than one channel, it was included only in the channel with the highest SNR, to prevent double counting. The analysis was performed blind without knowing the treatment group.

Neuronal units were excluded from further analysis if: 1) the amplitude of spike waveforms (defined by calculating peak to valley tick values (Plexon Offline sorter) versus time) decreased significantly before or during the injection to the detection limit or disappeared completely or temporarily into the noise, since this hampered an unambiguous evaluation of the drug effect, 2) the firing rate was < 0.4 Hz as very low firing rates also hampered the evaluation of the drug effect.

Different parameters such as firing rate of neuron, ISI, autocorrelation of spike times and the time taken for afterhyperpolarization phase (AHP) of the first derivative of an action potential were further analysed with custom written scripts in Matlab (version R2021b). The response of a single neuron to drug/vehicle injection as either excited, inhibited, or non-responding was determined by calculating the mean baseline firing rate of each neuron during a 2-minute period just before injection and comparing this to the mean firing rate during/after drug or vehicle injection (2-min interval: last minute during injection + first minute after injection). If the change in firing rate by drug or vehicle was 20% higher or lower than baseline, the effect was classified as excitatory or inhibitory, respectively. If the effect was smaller than 20% change, the neuron was termed non-responding i.e. no effect.

To determine the firing pattern of individual neurons, the same baseline period was analysed by calculating and plotting autocorrelation using the ‘xcorr’ and ‘coeff’ functions and ISI histograms of each neuron’s spiking times (bin size 20 ms). If the autocorrelation was 0 for lags from 0-200 ms, after which a clear peak appeared in both the autocorrelation and ISI distributions, the neuron was classified as regular firing (‘tonic regular’). When a bimodal distribution was observed at long lags in the autocorrelation or ISI distributions, the firing pattern of the neuron was defined as ‘slow oscillating’ where a peak at short lags corresponds to regular occurrence of spikes within a train and the second broader peak at long lags corresponds to the slow oscillating occurrence of these spike trains. If neither of these features was observed in the firing pattern of the neuron, the firing pattern was defined as irregular.

The afterhyperpolarization phase (AHP) of the extracellular action potential waveform was calculated as the full width at half maximum (FWHM) of the positive waveform peak using the Plexon offline sorter. The FWHM corresponds to the width of the spectral peak at half of its maximum amplitude. This was calculated as the distance between two points on the y-axis where the intensity of the curve is half of its peak value (see Fig. 5E).

### Statistical analysis

All statistical analysis was performed on raw data using SigmaPlot or GraphPad Prism software. The data distribution was tested with Shapiro–Wilk test and statistical tests for either normal or non-normal distributed data selected accordingly, e.g. student’s t-test or Wilcoxon rank sum test, respectively. When one-way ANOVA or Kruskal Wallis test for non-normal distributed data was used, p values of the post-hoc testing were corrected for multiple comparisons using the two-stage linear step-up procedure of Benjamini, Krieger and Yekutieli (Benjamini et al., 2006) that controls the false discovery rate (FDR) at the level q. All data are presented as mean ± SEM or median (interquartile range); p < 0.05 was considered statistically significant. In figures, the statistical significance levels are indicated by asterisks as follows: *p < 0.05, **p < 0.01, ***p < 0.001 and ****p < 0.0001.

## Acknowledgements

We thank Ada-Julia Kunnari and Rozalie Novakova for technical help. This study was financially supported by the Research Council of Finland (Research Fellowship (decision no 310765, 314104, 335396) to H.H.

