## Supplemental information for "Arginine vasopressin activates serotonergic neurons in the dorsal raphe nucleus during neonatal development *in vitro* and *in vivo*"

Figure S1.

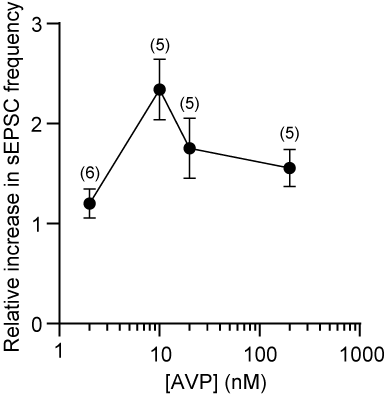

**Suppl Figure 1. AVP concentration response curve.** Normalized peak sEPSC frequency in response to 2, 10, 20 and 200 nM AVP. Numbers above the mean values indicate N= no of cells.

Figure S2

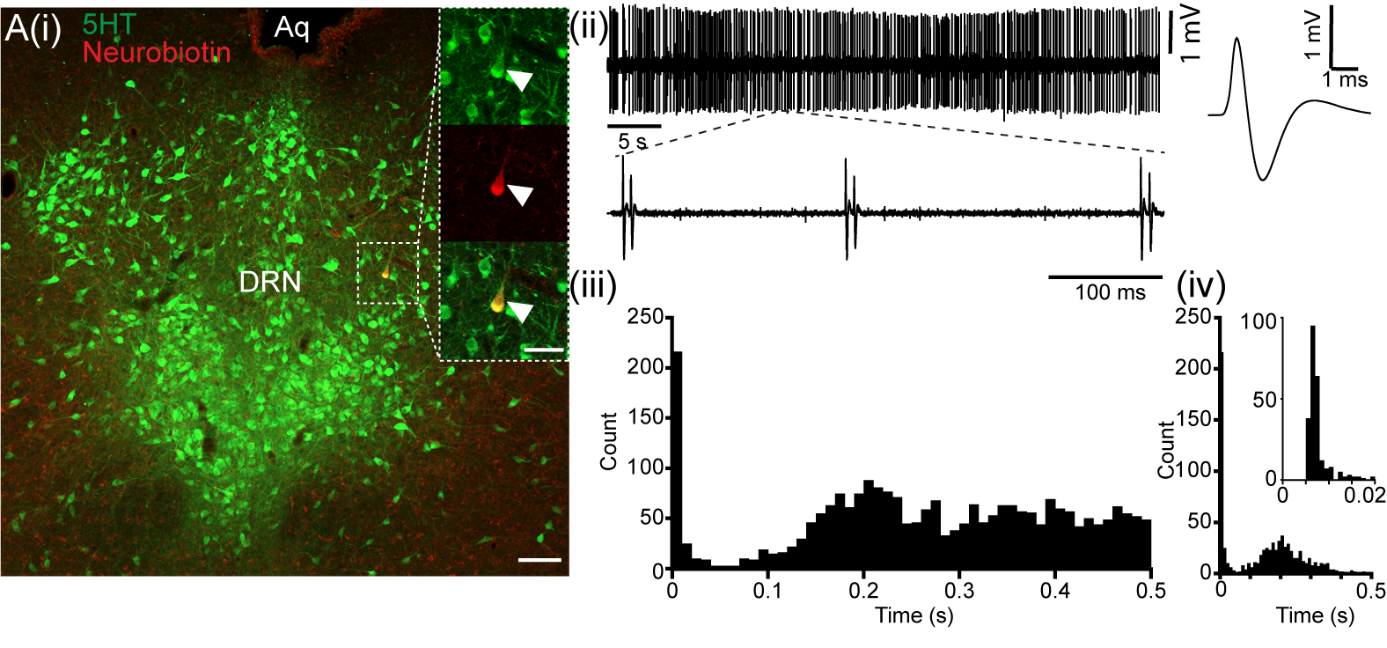

**Suppl Figure 2. Typical bursting 5-HT neuron firing doublets.** Photomicrograph **(i)** of 5-HT-positive neuron in neonatal dorsal raphe nucleus (DRN) (green, arrowhead) labelled with Neurobiotin (red, arrowhead) during *in vivo* electrophysiology. **(ii)** Corresponding spike train and waveform, **(iii)** autocorrelation histogram of spikes (bin size 10 ms) and **(iv)** inter-spike interval (ISI) histogram (bin size 10 ms) of the labelled neuron in **(i)**. Note the typical wide AP waveform and tonic regular firing of spike doublets (see enlarged trace) as characterised by peaks in the autocorrelation and ISI histogram at both short (see inset in ISI histogram, bin size=1 ms) and longer timescales.

Figure S3

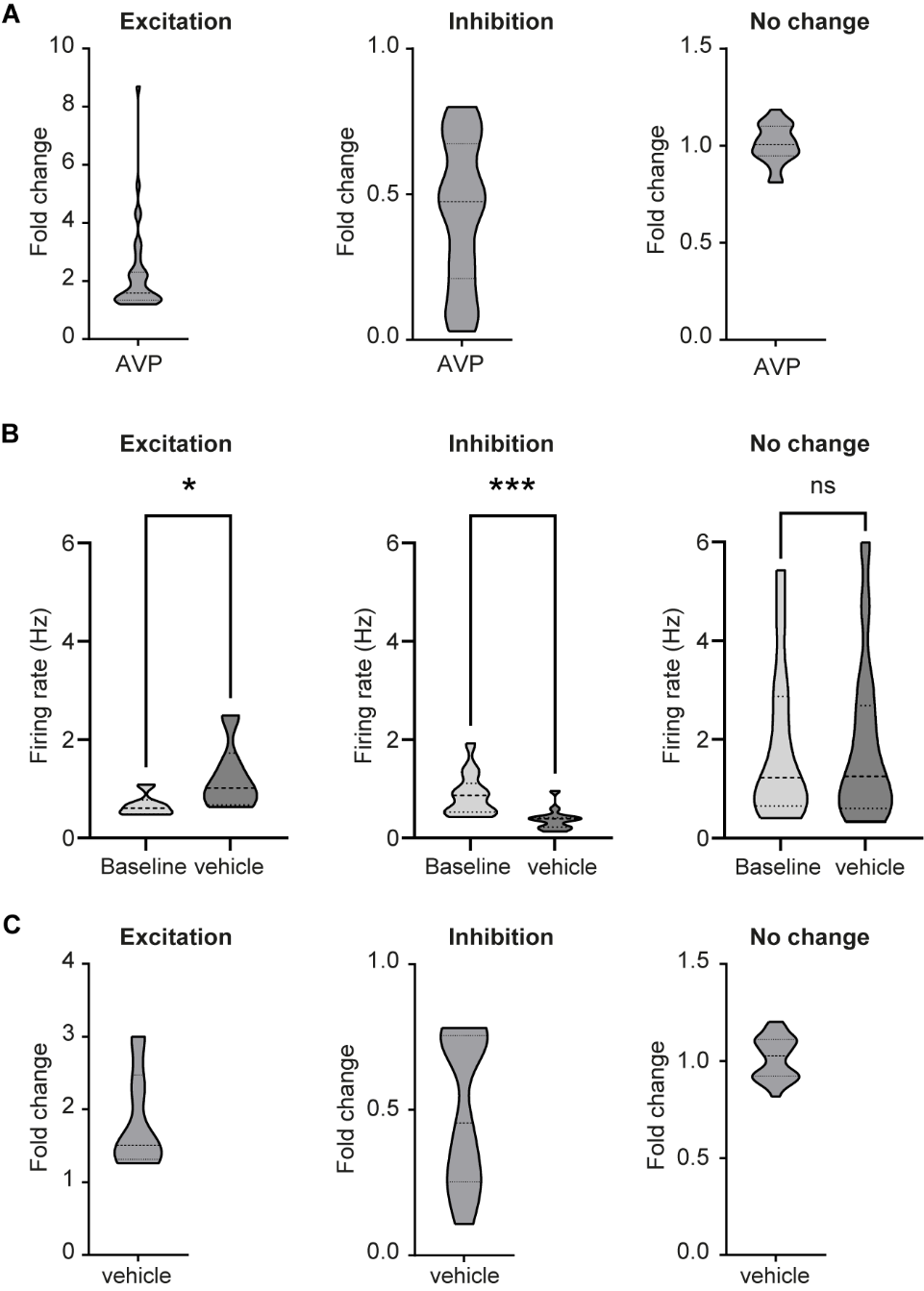

**Suppl Figure 3. Effect of vehicle and AVP on firing rates of DRN neurons *in vivo*. A,** Violin plots showing the fold change in firing rates by AVP of excited (N=36,left), inhibited (N=34, center) and non-reponding (N=35, right) DRN neurons after normalisation to own baseline firing rates. **B**, Violin plots of firing rates during baseline and after vehicle injection for excited (N=6, left), inhibited (N=13, center) and not responding (N=30, right) DRN neurons. **C**, Same as A for fold change in firing rates by vehicle injection. * P<0.05, *** P<0.001, paired Student’s t-test or Wilcoxon matched-pairs signed rank test.

Figure S4

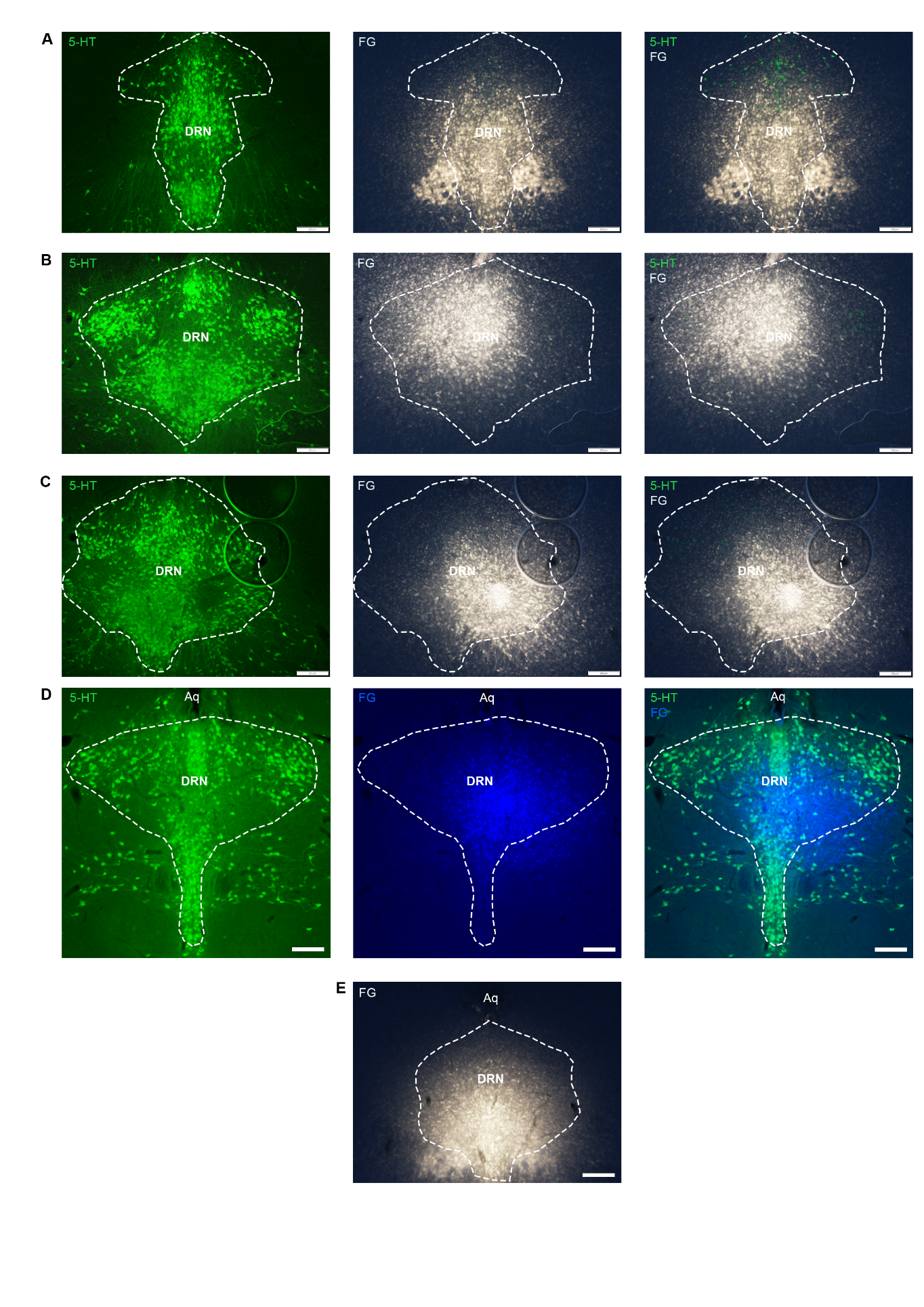

**Suppl Figure 4. Injection sites of FG are located in DRN. A-E,** Photomicrographs of coronal rat brain sections containing DRN showing the FG injection site for each pup (n=5) within DRN. Sections were processed for immunohistochemistry against 5-HT (green, A-D left) to estimate the outline of DRN in the sections (dashed line). The correct location of the FG injection site (white/blue, A-D middle) within the boundaries of DRN were confirmed (A-D, right). Sections from the pup shown in E were not processed for IHC against 5-HT. The outline of the DRN was estimated based on the size and location of the aqueduct (Aq) and the location of the FG injection site was confirmed as on the midline in ventral DRN. Scale bar: 200 µm.
